# Identification of key Y_4_R residues enables the discovery of selective non-peptide small-molecule agonists

**DOI:** 10.64898/2026.04.27.721169

**Authors:** Tim Pelczyk, Corinna Schüß, Mateusz Skłodowski, Fabian Liessmann, David Jordan, Vivian Ehrlich, Paul Eisenhuth, Jan Stichel, Albert O. Gattor, Max Keller, Jens Meiler, Annette G. Beck-Sickinger

## Abstract

G protein-coupled receptors (GPCRs) are central regulators of human physiology and disease, classifying them as relevant targets for therapeutic interventions. As transmembrane proteins, they convert extracellular signals into intracellular responses through agonist-induced conformational changes. Understanding how agonists stabilize active receptor conformations is decisive for rational drug design. In this study, we used the endogenous ligand pancreatic polypeptide (PP) and the cyclic hexapeptides UR-AK95c and UR-AK86c as molecular tools to determine key interactions critical for Y_4_R activation, which plays a crucial role in metabolic diseases. Guided by molecular docking, we systematically replaced Y_4_R residues and assessed activation. The *in vitro* and *in silico* studies delineated a key Y_4_R activation interface centered around the conserved C-terminal RXRY-NH_2_ motif of the peptides and, separately, identified receptor residues with distinct peptide-specific functional effects. Next, we performed an ultra-large library screening (ULLS) and experimentally validated three predicted hits as selective Y_4_R agonists that engage in a substantial subset of the identified critical receptor contacts. This study demonstrates how GPCR activation interface knowledge can be translated into the discovery of novel small-molecule agonists and outlines a general strategy for advanced GPCR drug discovery.

## Introduction

G protein-coupled receptors (GPCRs) constitute the largest superfamily of transmembrane proteins in humans and play a central role in cellular signal transduction by converting extracellular stimuli into intracellular responses (***Latorraca et al., 2017***). Ligand binding stabilizes distinct receptor conformations, shifting GPCRs from inactive to active states and thereby initiating downstream signaling events primarily mediated by heterotrimeric G proteins and arrestins (***Filipek, 2019***). Due to their importance in the regulation of physiological processes and excellent drugability, GPCRs represent one of the most relevant classes of therapeutic targets in modern pharmacology (***Hauser et al., 2017***).

Structure-based drug design has emerged as a powerful strategy for the development of GPCR-targeting drugs. Advances in cryo-electron microscopy (cryo-EM) and X-ray crystallography have led to a substantial increase in available GPCR structures (***Zhang et al., 2024***), with more than 1000 inactive and active structures reported by 2023 (***Liu et al., 2024***). Despite this rich structure collection, receptor models often fail to precisely reflect ligand-receptor interactions that are crucial to stabilize specific signaling-competent conformations as only neighborhood is considered. Defining the activation interfaces is therefore important for the successful rational design of GPCR agonists (***Mason et al., 2012***).

The neuropeptide Y Y_4_ receptor (Y_4_R) is a particularly interesting yet underexploited GPCR target (***Schüß et al., 2024***; ***Zhu et al., 2023***). Y_4_R is a member of the neuropeptide Y system, a multireceptor/multiligand family that comprises three additional neuropeptide Y receptor subtypes, the Y_1_R, Y_2_R_,_ and Y_5_R, and three C-terminally amidated 36-amino acid peptide ligands, neuropeptide Y (NPY), peptide YY (PYY), and pancreatic polypeptide (PP) (***Lindner et al., 2008***). While Y_1_R, Y_2_R, and Y_5_R show a preference for NPY and PYY, the Y_4_R is predominantly activated by PP (***Pedragosa-Badia et al., 2013***), an endocrine hormone that is excreted postprandially by PP-cells of the pancreatic islets (***Lonovics et al., 1981***). Y_4_R is primarily expressed in the periphery, with highest abundance in the colon (***Cox and Tough, 2002***), small intestine (***Goumain et al., 1998***), and pancreas (***Kim et al., 2014***). To a lower amount, it is additionally localized in specific regions of the central nervous system (***Campbell et al., 2003***; ***Parker and Herzog, 1999***) characterized by an incomplete blood-brain barrier (***Schüß et al., 2024***). Functionally, Y_4_R suppresses intestinal motility (***Moriya et al., 2010***) and secretion (***Tough et al., 2006***), modulates insulin levels by somatostatin (***Kim et al., 2014***), and has been associated with β-cell protection (***Khan et al., 2017***). In the hypothalamus and brainstem, Y_4_R signaling contributes to the regulation of appetite suppression (***Sainsbury et al., 2010***) and energy expenditure (***Holzer et al., 2012***). This combination of peripheral accessibility and metabolic control highlights Y_4_R agonism as an attractive therapeutic strategy for the treatment of obesity and metabolic disorders (***Yulyaningsih et al., 2011***; ***Zhu et al., 2023***). Nevertheless, the lack of clinically approved Y_4_R-targeting drugs underscores the need for new ligand discovery approaches.

In this study, three equipotent Y_4_R peptide agonists, PP and the cyclic hexapeptides UR-AK95c and UR-AK86c (***Konieczny et al., 2021***), were used as molecular tools to determine the key interface required for receptor activation. Based on the PP-Y_4_R-G_i1_ cryo-EM structure (PDB:7X9C) (***Tang et al., 2022***), we performed molecular docking of these peptides into the orthosteric receptor site, which systematically guided subsequent mutagenesis studies. This approach revealed both shared and peptide-specific receptor interactions that delineate a key activation interface stabilizing the active Y_4_R conformation. In addition, functional analysis of truncated UR-AK86c variants (***Konieczny et al., 2020***) revealed that multiple structural elements of the hexapeptides contribute to their high potency, while the C-terminal tetrapeptide is sufficient to maintain nanomolar activity. The insights into the Y_4_R activation interface subsequently allowed for an ultra-large library screening (ULLS) that identified small-molecule agonist candidates, three of which were experimentally validated as subtype-selective Y_4_R agonist hits. Together, these findings outline a broadly applicable strategy for translating knowledge of critical receptor interactions into GPCR agonist discovery.

## Results

Functional assays validate UR-AK95c and UR-AK86c as suitable molecular tools to investigate molecular mechanisms underlying Y_4_R activation UR-AK95c and UR-AK86c bind to Y_4_R with picomolar affinity by mimicking the C-terminal pentapeptide of PP (Figure 1A) (***Konieczny et al., 2021***), which was shown to bind deeply in the helical bundle of Y_4_R (***Tang et al., 2022***). To confirm their value as tools for probing Y_4_R, we compared their ability to induce G protein signaling with the endogenous ligand PP. In a BRET-based G_i3_ protein assay, which directly quantifies the dissociation of the heterotrimeric G protein (***Schihada et al., 2021***), PP showed an half maximal effective concentration (EC_50_) value of 0.71 nM (pEC_50_ 9.15 ± 0.09) at Y_4_R, whereas the hexapeptides display a slightly higher potency with EC_50_ values of 0.11 nM (pEC_50_ 9.96 ± 0.11) and 0.45 nM (pEC_50_ 9.49 ± 0.15) for UR-AK95c and UR-AK86c, respectively (Figure 1B, Figure 1 -table supplement 1). A kinetic set-up of this assay demonstrated a faster decline of netBRET following stimulation with the cyclic hexapeptides over time compared to PP, indicating a more rapid dissociation of the heterotrimeric G protein, and thus a faster receptor activation (Figure 1C). In an IP-one accumulation assay, an equilibrium assay measuring the G_q_ protein-dependent inositol-monophosphate (IP-one) accumulation (***Besserer-Offroy et al., 2020***), PP demonstrated a slightly higher Y_4_R activity compared to the hexapeptides with EC_50_ values of 0.16 nM (pEC_50_ 9.79 ± 0.06) for PP, 0.29 nM (pEC_50_ 9.53 ± 0.12) for UR-AK95c, and 0.41 nM (pEC_50_ 9.39 ± 0.09) for UR-AK86c (Figure 1D, Figure 1 -table supplement 1). As Y_4_R endogenously couples to G_i/o_ proteins, the assay was adapted to Y_4_R by using the chimeric G protein Δ6Gα_qi4myr_ (Kostenis, 2001). To assess whether the improved binding selectivity of UR-AK95c and UR-AK86c compared to PP also translates to functional selectivity, the three Y_4_R high-affinity peptides (and NPY) were tested in IP-one accumulation assays at all neuropeptide Y receptor subtypes. Here, PP showed full agonistic activity at all Y receptor subtypes with potencies of 42.8 nM (pEC_50_ 7.37 ± 0.08) at Y_1_R, 347 nM (pEC_50_ 6.46 ± 0.15) at Y_2_R, and 8.14 nM (pEC_50_ 8.09 ± 0.05) at Y_5_R, whereas the hexapeptides displayed significantly lower activities at Y_1_R, Y_2_R, and Y_5_R. Interestingly, at Y_2_R and Y_5_R, UR-AK86c, which contains Tyr on position 2, seems to be less potent compared to UR-AK95c, which features a Trp at the corresponding position (Figure 1A, D). In addition, the effect of the allosteric antagonists *(S)*-VU0637120 (***Schüß et al., 2021***) and the positive allosteric modulator VU0506013 (***Schüß et al., 2023***) was investigated on the peptide-induced Y_4_R activation using Ca^2+^-flux assays. Interestingly, the activity of all three peptides was modulated by both compounds to a similar degree with an approximately 10-fold reduction of activity with *(S)*-VU0637120 and a roughly 3-fold increased potency with VU0506013, suggesting a similar binding mode of the peptides at Y_4_R (Figure 1E, Figure 1 -table supplement 2).

**Figure 1:**
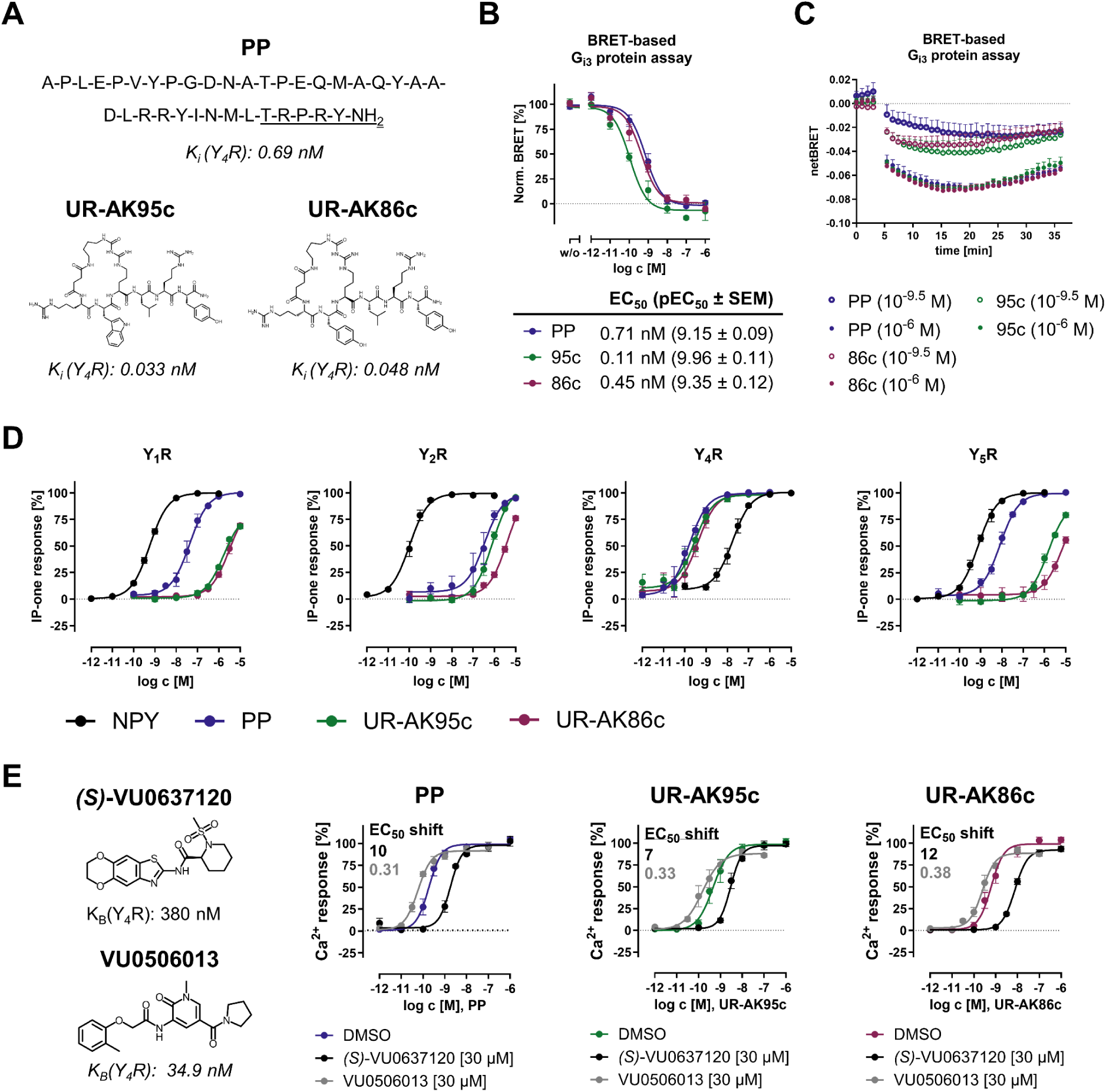
Functional studies of PP, UR-AK95c, and UR-AK86c. A) Sequence of PP and structure of UR-AK95c and UR-AK86c. The C-terminal pentapeptide of PP mimicked by the hexapeptides is underlined. Ki values were previously reported (***Konieczny et al., 2021***). B) Concentration-response curves, EC50, and pEC50 value of PP, UR-AK95c, and UR-AK86c at Y4R obtained by BRET-based Gi3 protein assay performed in COS-7 cells transiently transfected with untagged Y_4_R(wt) and Gi3 protein-based tricistronic activity sensor (Gi3-CASE). C) Kinetic measurement of ligand-induced G-protein dissociation over time obtained by BRET-based Gi3 protein assay. D) Concentration-response curves of NPY, PP, UR-AK95c, and UR-AK86c at human neuropeptide Y receptor subtypes obtained by IP-one accumulation assays in stably transfected COS-7_Y1/2/4/5R_ GαΔ6qi4myr cells. E) Structure of Y4R allosteric antagonist *(S)*-VU0637120 and Y_4_R positive allosteric modulator VU0506013 and concentration-response curves of PP, UR-AK95c, and UR-AK86c in presence of DMSO, 30 µM of *(S)*-VU0637120, or 30 µM of VU0506013. Data were obtained by Ca^2+^-flux assay in COS-7 cells stably transfected with C-terminally eYFP-tagged Y4R and GαΔ6qi4myr. All data are presented as the mean ± SEM from at least three independent experiments, each performed in triplicates.

### Docking reveals conserved and peptide-specific Y_4_R interactions

To elucidate how the hexapeptides mimic the activity of the endogenous ligand in the orthosteric site, initial docking models of PP, UR-AK95c, and UR-AK86c were generated using Rosetta, supported by RDKit (***Landrum et al., 2025***) and BCL (***Brown et al., 2022***), based on the PP-Y_4_R-G_i1_ cryo-EM structure (***Tang et al., 2022***). Subsequent mutagenesis results guided iterative refinement of these models. The top-ranking models, based on their agreement with the mutagenesis data, are shown in Figure 2 and were used as a basis for per-residue energy breakdown (Figure 2-figure supplements 1-3). The molecular docking revealed an overlapping binding mode of the conserved C-terminal -RY-NH_2_ motif deep in the orthosteric Y_4_R pocket, with PP located slightly deeper in the helical receptor bundle. According to the docking model, the C-terminal amide of the peptides is analogously positioned in proximity to T2.61, Q3.32, and H7.39 of Y_4_R, whereas the side chains of Y^36^/Y^6^ are predicted to form hydrophobic and polar contacts with several residues, including C2.57, C3.33, V3.36, F4.60, L5.42, L6.51, and Q5.46, respectively. In addition, the R^35^/R^5^ side chains of the peptides are located similarly near E203^ECL2^, N6.55, and D6.59. This conserved binding mode of the C terminus of the peptides is also supported by the per-residue energy breakdown, which indicates favorable (negative) energy scores between the - RY-NH_2_ motif of the peptides and the aforementioned residues, with the exception of PP at positions T2.61 and N6.55. In contrast, for P^34^ of PP and the corresponding L^4^ of the hexapeptides, the docking model revealed some differences in the binding mode. Whereas all three peptides are in proximity to Y2.64 and W107^ECL1^ according to the interface analysis, L^4^ of the hexapeptides expands towards TM3 and the top of TM4, resulting in additional contacts with C3.25, S3.28, A3.29, and I179^ECL2^. In addition, the R^33^/R^3^ side chains of PP and the hexapeptides are located similarly in proximity to E6.58 and F7.35. However, R^33^ of PP forms additional contacts with F6.54 and N7.32, whereas, according to the per-residue energy breakdown of UR-AK86c, the carbamoylated R^3^ could partially mimic contacts of the α-helix of PP with ECL3. Interestingly, with the exception of weak contacts with C7.29, the succinyl linker, which connects the carbamoylated R^3^ with the N terminus of the hexapeptides, does not substantially engage in contacts with Y_4_R. Moreover, the binding mode of the analogous residues T^32^ of PP and W^2^/Y^2^ of the hexapeptides reveal the most pronounced differences in the binding between the Y_4_R peptides. Whereas the docking shows prominent contacts of T^32^ of PP with D2.68 and W107^ECL1^, W^2^ and Y^2^ of the hexapeptides predominantly interact with ECL2 residues, including N182^ECL2^, V183^ECL2^, F184^ECL2^, V200^ECL2^ and T202^ECL2^. In contrast, these positions are involved in interactions with PP through residues located in its α-helix. The binding mode of R^1^ of the hexapeptide is less defined by the docking models, however contacts with E203^ECL2^ of Y_4_R are possible.

**Figure 2:**
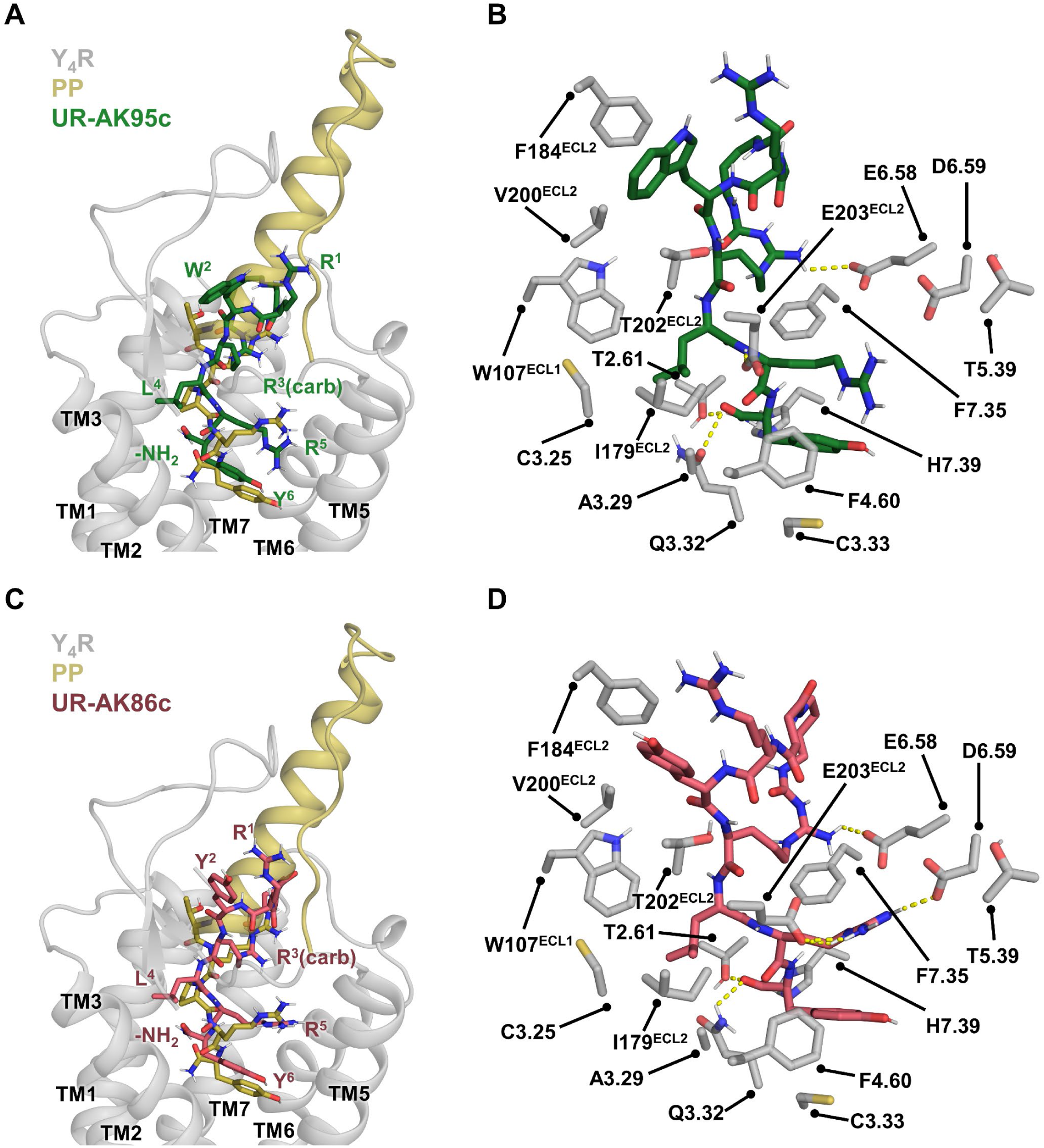
Docking models of UR-AK95c and UR-AK86c at Y4R compared to PP. A) and C) show the overall binding modes of UR-AK95c (green, A) or UR-AK86c (red, C) superimposed with PP (yellow, A and C) at Y4R, generated using Rosetta based on the PP-Y_4_R-G_i1_ cryo-EM structure (PDB:7X9C). TM4 is omitted for better visualization. B) and D) highlight receptor residues in proximity to UR-AK95c (green, B) or UR-AK86c (red, D) that contribute < −1 Rosetta Energy Units (REU) to the interaction energy with at least one of the hexapeptides, as determined by a per-residue energy breakdown. The corresponding contact maps of this interface analysis for PP, UR-AK95c, and UR-AK86c are provided in Figure 2-figure supplement 1-3.

### Mutagenesis studies identify Y_4_R residues relevant for PP, UR-AK95c, and UR-AK86c

As molecular docking suggests neighborhood of residues, *in vitro* mutagenesis studies, followed by activity assays is required to identify functionally relevant contacts. Residues of human Y_4_R were selected based on the preliminary docking models and a previously published MD simulation of UR-AK95c and UR-AK86c at a κ-opioid receptor-based Y_4_R homology model (***Konieczny et al., 2021***). In total, 38 residues were substituted to alanine and the respective receptor variants were analyzed in IP-one accumulation assay with PP, UR-AK95c, and UR-AK86c. The corresponding results are summarized in Figure 3 and Figure 3 -table supplement 1.

**Figure 3:**
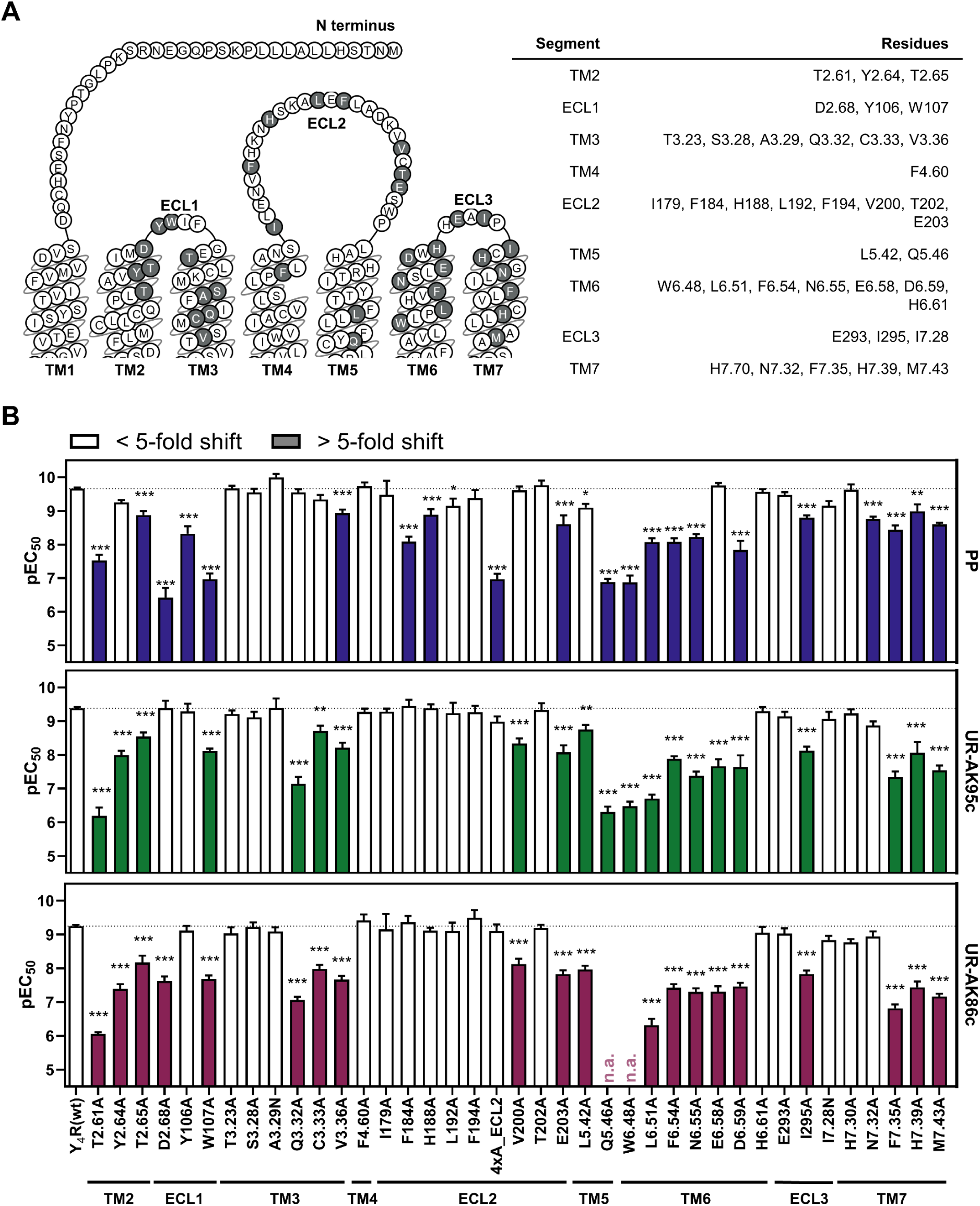
Mutagenesis studies of PP, UR-AK95c and UR-AK86c at Y4R. A) Y_4_R snake plot. Residues selected for mutagenesis studies are marked in grey. Residues are named according to Ballesteros-Weinstein nomenclature (***Ballesteros and Weinstein, 1995***). Snake plot was adapted from GPCRdb.org (***Pándy-Szekeres et al., 2018***). B) pEC_50_ values of PP, UR-AK95c, and UR-AK86c determined for Y_4_R single- and multiple residue variants. Mutagenesis data were obtained by IP-one accumulation assay conducted in COS-7 cells transiently transfected with C-terminally eYFP-tagged Y4R variant and Δ6Gαqi4myr. Significance was determined by one-way ANOVA with Dunnett’s post-test. *, P < 0.033, **, P < 0.002, ***, P < 0.001. 4xA_ECL2, Y_4_(F184A, H188A, L192A, F194A). Data represent the mean ± SEM from at least three experiments, each performed in triplicates.

For PP, the effects of the Y_4_R substitutions on receptor signaling, which were previously reported as part of the PP-Y_4_R-G_i1_ cryo-EM structure (PDB:7X9C) (***Tang et al., 2022***), have been confirmed. In addition, newly generated Y_4_R residue variants have been identified with distinct effects on PP: While T2.65A, Y106^ECL1^A, W107^ECL1^A, 4xA_ECL2, and W6.48A significantly reduced PP activity, the substitutions T3.23A, S3.28A, A3.29N, I179A, F194A, V200A, H6.61A, H7.30A, and I297N had no significant effect on PP-induced receptor signaling.

To compare the significance of these contacts between the hexapeptides and PP, all Y_4_R residue variants were characterized with UR-AK95c and UR-AK86c, distinguishing between conserved and peptide-specific functional effects. In TM2, substitutions at T2.61, Y2.64, and T2.65 significantly reduced the activity of both hexapeptides, while D2.68 selectively induced a 42-fold EC_50_ shift for UR-AK86c compared to wildtype Y_4_R, suggesting differences in the binding mode of the hexapeptides at this receptor position. In ECL1, Y106^ECL1^ had no impact, while replacement of W107^ECL1^ with alanine resulted in a loss in potency for both UR-AK95c and UR-AK86c. In TM3 and TM4, the residues close to the extracellular side failed to alter peptide potency, whereas substitutions in the TM region, including Q3.32, C3.33, and V3.36, impaired peptide activity, suggesting that the hexapeptides engage TM3 relatively deep in the helical bundle. Except for V200^ECL2^ and E203^ECL2^, residues in ECL2 did not significantly affect hexapeptide-mediated receptor signaling, disfavoring a major contribution of this extracellular segment to receptor activation. In contrast, substitutions of several residues in TM5 and TM6, including Q5.46, L6.51, F6.54, N6.55, and D6.59, to alanine resulted in a substantially reduced activity of both hexapeptides, indicating an important role of TM5 and TM6 for the hexapeptide-induced receptor signaling. In ECL3 and the upper region of TM7, only the I295^ECL3^A variant showed a significant loss of potency for the hexapeptides. All receptor variants located deeper in TM7, including F7.35, H7.39, and M7.43 resulted in a reduced activity of for UR-AK95c and UR-AK86c, highlighting the relevance of these positions for the hexapeptides.

As most of the differences between the hexapeptides and PP were observed in the extracellularly located receptor segments, ECLs of Y_4_R were exchanged with those of the phylogenetically closely related (***Larhammar and Salaneck, 2004***) Y_1_R. This resulted in three Y_4_/Y_1_R loop chimeras (***Schüß et al., 2021***), which were tested with PP, UR-AK95c, and UR-AK86c. The data from these chimera studies are summarized in Figure 3-figure supplement 1 and Figure 3-table supplement 1. While PP retained wildtype-like potency at all Y_4_/Y_1_R loop chimeras, UR-AK95c and UR-AK86c were differentially influenced by the loop exchanges. At Y_4_(ECL1_Y_1_), only UR-AK86c exhibited a significant loss in potency, whereas exchanging ECL2 of Y_4_R to that of Y_1_R resulted in a modest but significant gain-of-function for the hexapeptides. Notably, Y_4_(ECL3_Y_1_) induced a loss of activity for both UR-AK95c and UR-AK86c, which suggests ECL3 as a potential selectivity driver that mediates the improved selectivity of both hexapeptides for Y_4_R towards the phylogenetically related Y_1_R.

### Mutagenesis and molecular docking correlate and delineate the key Y_4_R activation interface

Functional studies showed that the truncated peptides UR-AK95c and UR-AK86c activate Y_4_R with potencies comparable to that of the full-length ligand, indicating that Y_4_R activation can be achieved by rather small peptides (Figure 1). This is further emphasized by the docking models, revealing that UR-AK95c and UR-AK86c form substantially smaller receptor interfaces, measuring approximately 1200 Å^2^ respectively, whereas PP extends further towards the extracellular loops, resulting in an estimated interface area of 2800 Å^2^ (Figure 4A). The surface area was estimated by subtracting the solvent accessible surface area of the peptide-Y_4_R-complex from the sum of the solvent accessible interface area of Y_4_R and the peptide using PyMol (***DeLano, 2026***).

**Figure 4:**
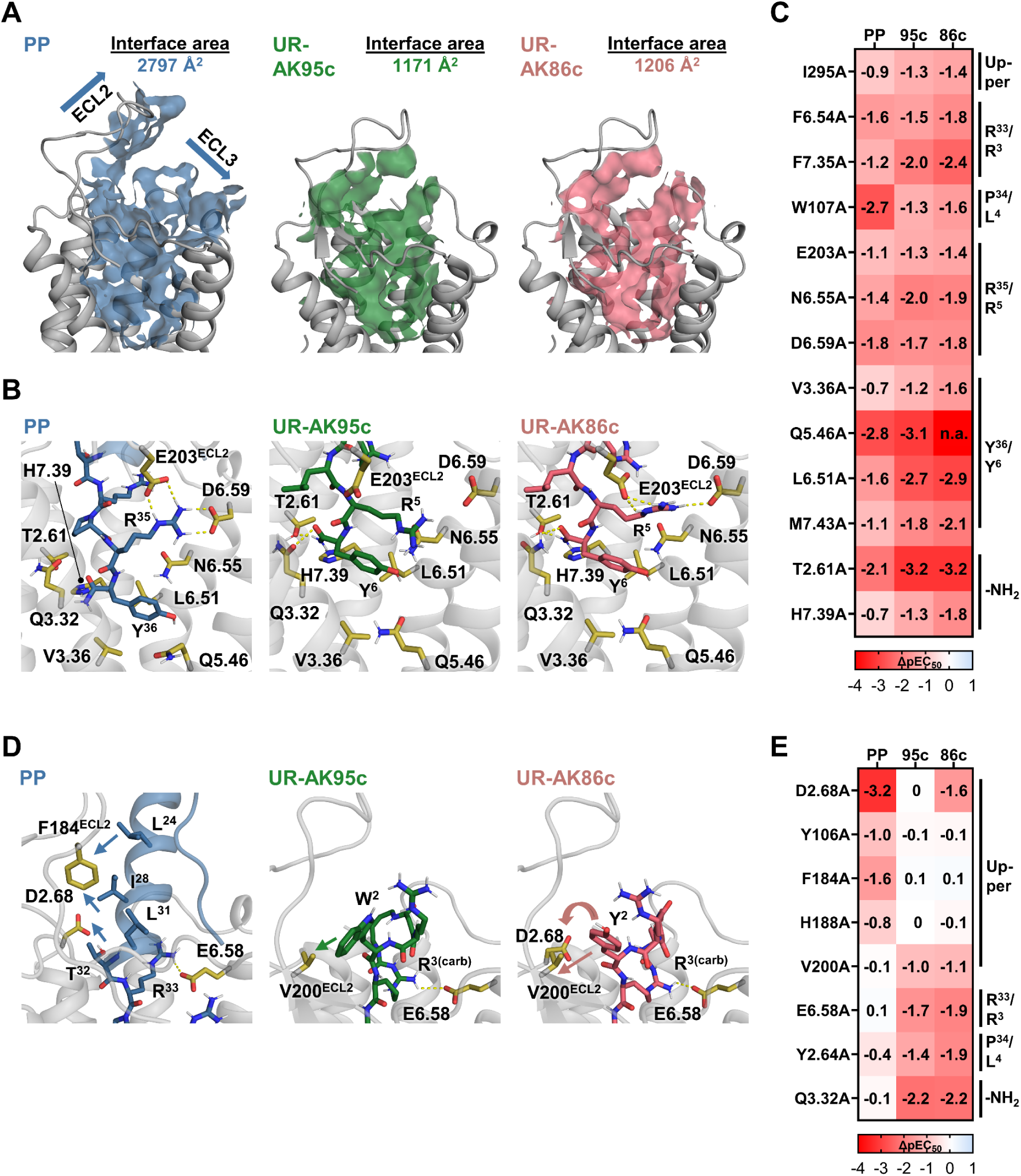
Key Y4R activation interface and receptor residues mediating peptide-specific interactions. A) Area of Y_4_R within a distance of 5 Å of PP (blue), UR-AK95c (green), or UR-AK86c (red). Surface area was estimated by subtracting the solvent accessible surface area of the peptide-Y_4_R-complex from the sum of the solvent accessible interface area of Y_4_R and the peptide using PyMol (***DeLano, 2026***). B) Selected Y_4_R key residues interacting with the peptides’ conserved C-terminal -RY-NH_2_ motif. C) ΔpEC_50_ values of Y_4_R variants displaying more than 5-fold reduction in potency across all three peptides compared to Y_4_R(wt). D) Selected Y_4_R residues that contribute differentially to the interactions with the three peptide agonists. E) ΔpEC_50_ values of Y_4_R variants displaying pronounced differences between the peptides. For better visualization, TM4 is omitted in all structural snapshots.

To verify the correlation between the *in silico* and *in vitro* data, mutagenesis results were compared with the molecular docking of the peptides at Y_4_R. Mutagenesis studies identified 15 Y_4_R residues that induced a ΔpEC_50_ ≥ 0.7 for all peptides, corresponding to an at least 5-fold reduction in potency compared to wildtype receptor. The per-residue energy breakdown showed that 13 of these residues contribute to peptide binding at Y_4_R (Figure 2-figure supplements 1-3). These residues were identified as the key Y_4_R interface (Figure 4C). Notably, nine of these residues are predicted to form conserved interactions with the C-terminal -RY-NH_2_ motif shared by all three peptides, with T2.61, V3.36, Q5.46, L6.51, H7.39, and M7.43 interacting with the tyrosine amide and E203^ECL2^, N6.55, and D6.59 interacting with R^35^/R^5^ of the peptides (Figure 4B). This indicates a major contribution of this peptide segment to Y_4_R activity. The docking studies further suggest that peptide positions P^34^/L^4^ and R^33^/R^3^ form analogous contacts with W107^ECL1^, F6.54, and F7.35, respectively. Interestingly, mutagenesis showed that I295^ECL3^, which is engaged by R^26^ and M^30^ in PP, is also relevant for the hexapeptides. This effect could be attributed to interactions with the macrocycle in the hexapeptides, as shown by the interface analysis of UR-AK86c (Figure 2 -table supplement 1-3). In addition, mutagenesis studies determined Y^4^R residues with distinct effects on the peptide activity that are primarily located in the extracellular receptor regions and largely consistent with the docking models (Figure 4E). Y106^ECL1^ and W107^ECL1^ are more relevant for PP, likely through contacts with T^32^ of PP, which is replaced by W^2^ or Y^2^ in the hexapeptides. Interestingly, D2.68 also contributes to UR-AK86c activity, indicating transient or water-mediated interactions with Y^2^ that are not captured by the molecular docking. Within ECL2, F184^ECL2^ and H188^ECL2^ are more relevant for PP, while V200^ECL2^ is crucial for the hexapeptides. The molecular docking shows that L^24^, I^28^, and L^31^ in PP are more orientated towards the respective ECL2 residues, whereas W^2^ and Y^2^ of the hexapeptides engage ECL2 through contacts with V200^ECL2^. In addition, Y2.64 and E6.58 are more important for the hexapeptides. Docking suggests that this effect may arise from the P^34^ to L^4^ substitution in the hexapeptides and the cyclization-induced conformational restriction of the R^3^ sidechain, respectively. Notably, Q3.32 shows higher functional relevance for the hexapeptides than for PP, although docking predicts conserved interactions with the C-terminal amide across all peptides.

### Structure-activity insights from truncated UR-AK86c derivatives

To elucidate the critical receptor activation interface at the peptide site, three truncated UR-AK86c analogues were evaluated in IP-one accumulation assay at Y_4_R. These peptide analogues included [Agp^5^]-UR-AK86c, containing 2-amino-3-guanidino-propionic acid (Agp) at position 5 instead of Arg, the linear precursor peptide RYRLRY-NH_2_, and the N-terminally truncated tetrapeptide RLRY-NH_2_ (***Konieczny et al., 2020***) (Figure 5A). The results are summarized in Figure 5 and Figure 5 -table supplement 1. All peptide variants demonstrated a reduced potency at Y_4_R compared to UR-AK86c with EC_50_ values of 5.46 nM, 9.00 nM, and 16.7 nM. Thus, all tested structural components contribute to the high activity of the cyclic hexapeptides at Y_4_R, however a linear tetrapeptide is sufficient to retain nanomolar potency. To investigate the contribution of the individual peptide components to selectivity, all peptides were also tested at the other neuropeptide Y receptor subtypes. None of the peptides induced maximal receptor activation within the tested concentration range. Substitution of R^5^ to Agp^5^ in UR-AK86c abolished activity at Y_1_R and Y_2_R, while Y_5_R activity was unaffected. In contrast to Y_4_R, deletion of the cyclic structure did not reduce peptide activity at Y_1_R, Y_2_R, and Y_5_R, underscoring the significance of the cyclic structure for enhanced Y_4_R activity and selectivity. Notably, the tetrapeptide RLRY-NH_2_ was inactive at Y_1_R and Y_5_R but displayed higher potency than RYRLRY-NH_2_ and UR-AK86c at Y_2_R, suggesting an important role of the N-terminal RY motif in Y_2_R/Y_4_R discrimination.

**Figure 5:**
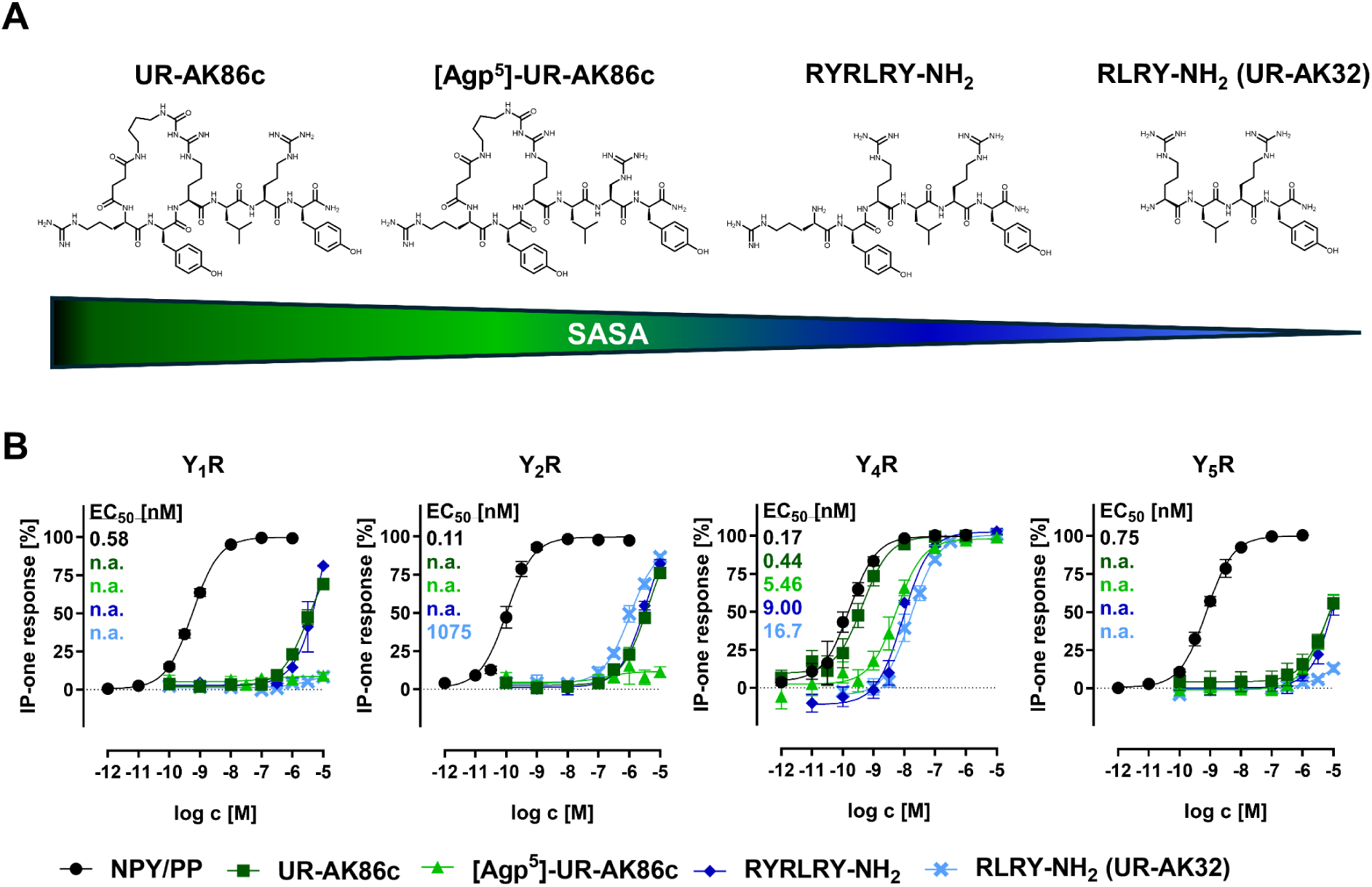
Functional studies of UR-AK86c and truncated derivatives at neuropeptide Y receptor subtypes. A) Structure of UR-AK86c, [Agp^5^]-UR-AK86c, the linear hexapeptide RYRLRY-NH2, and UR-AK32 (***Konieczny et al., 2020***) in descending order of solvent-accessible surface area (SASA). B) Concentration-response curves of PP at Y4R; NPY at Y1R, Y2R, and Y5R, and UR-AK86c, [Agp^5^]-UR-AK86c, RYRLRY-NH2, and UR-AK32 at all neuropeptide Y receptor subtypes obtained through IP-one accumulation assay conducted in stably transfected COS-7_Y1/2/4/5R_GαΔ6qi4myr cells. All data are presented as the mean ± SEM from at least three independent experiments, each performed in triplicates.

### ULLS translates Y_4_R activation hotspots into hits for small-molecule agonists

In the next step, insights into the receptor activation interface and peptide pharmacophore informed an ULLS for small-molecule Y_4_R agonists. The screening was conducted using Enamine’s REAL (Readily AccesibLe) database, which comprised more than 36 billion synthetically accessible virtual compounds at the time of this study. From this collection, more than 560,000 compounds were evaluated against the Y_4_R structure using the Rosetta Evolutionary Ligand (REvoLd) framework (***Eisenhuth et al., 2025***). In addition to standard scoring and physicochemical filtering, candidate compounds were prioritized based on their predicted engagement of Y_4_R residues identified in this study as critical for activity transmission. Following this rigorous multistep filtering strategy, 53 compounds were selected and ordered in three successive batches for experimental validation in an IP-one accumulation assay (Figure 6 -table supplement 1). Three compounds showed significant Y_4_R activity at 100 µM, with 70 % (Z9407529782), 56 % (Z8878907875), and 45 % (Z9239258318) of maximal PP-response, respectively (Figure 6A). These compounds were counter-screened at the remaining neuropeptide Y receptor subtypes, where none of the compounds induced significant IP-one accumulation, indicating a Y_4_R-dependent and subtype-selective activity (Figure 6B).

**Figure 6:**
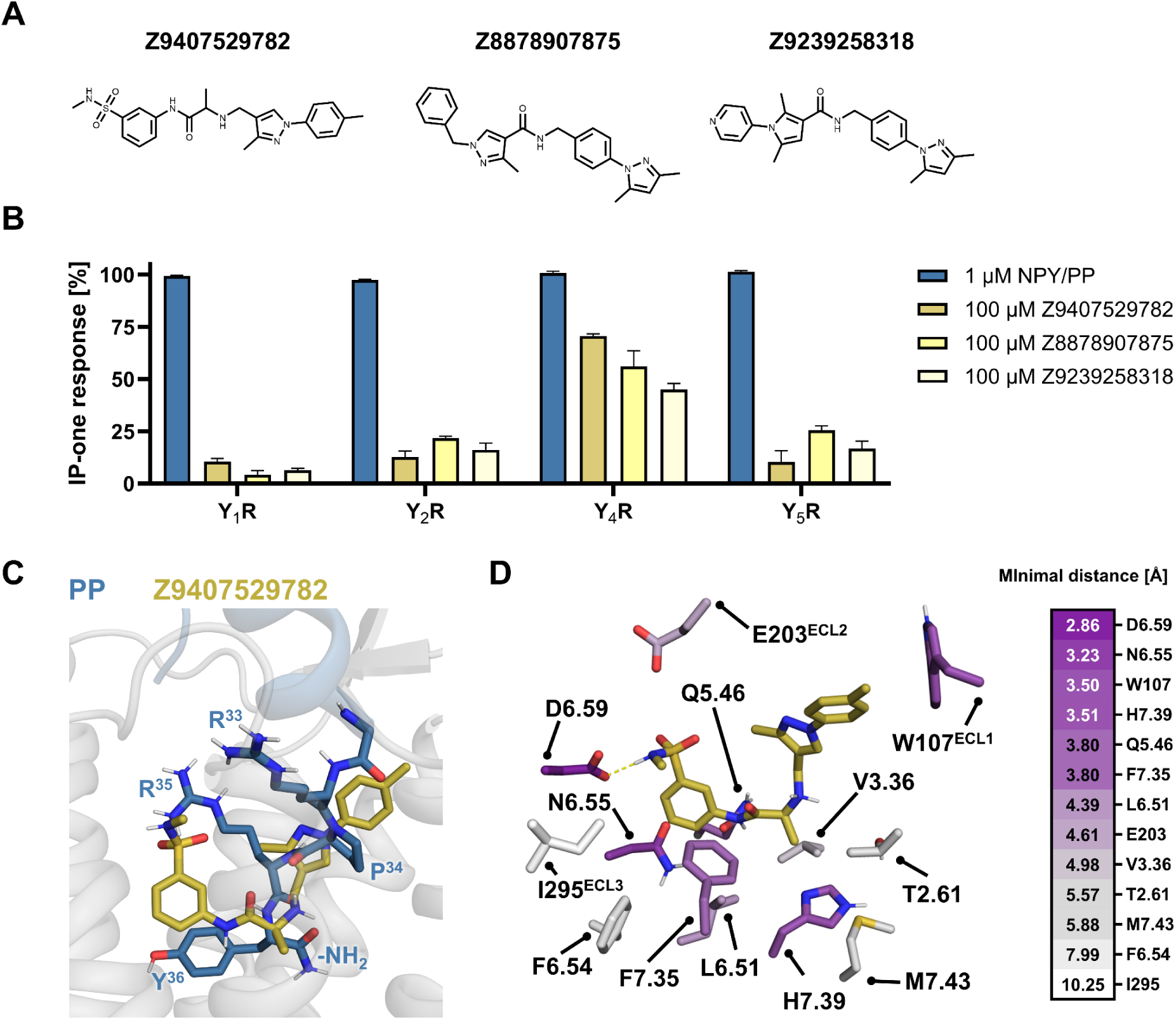
Identification of hits from ULLS. A) Structures of Z8878907875, Z9239258318, and Z9407529782. B) Activation of neuropeptide Y receptor subtypes by 100 µM Z8878907875, 100 µM Z9239258318, and 100 µM Z9407529782 in comparison with 1 µM PP or 1 µM NPY, respectively. Data were obtained through IP-one accumulation assay conducted in stably transfected COS-7_Y1/2/4/5R_ GαΔ6qi4myr cells. All data are presented as the mean ± SEM from at least three independent experiments, each performed in triplicates. C) Overlay of PP and Z9407529782 in the Y4R binding pocket as predicted by FlexPepDock and RosettaLigand, respectively. D) Representative docking pose of Z9407529782 at Y4R, with receptor residues shown as sticks, which were defined as key Y4R residues in this study. Residues are colored according to their minimal distance to Z9407529782.

The most potent and selective hit Z9407529782 was redocked at Y_4_R and subjected to tethered minimization using RosettaLigand (***Lemmon and Meiler, 2012***) and Molecular Operating Environment (MOE) (***Chemical Computing Group ULC, 2026***) respectively. Interestingly, the resulting docking model revealed an overlapping binding mode of Z9407529782 and the C-terminal residues of PP, suggesting a similar mechanism of receptor activation. While the sulfonamide of the benzene sulfonyl group of Z9407529782 overlaps with the R^35^ sidechain of PP, the benzene group potentially mimics the aromatic Y^36^ ring. In addition, the alanine-derived amide linker of Z9407529782 is similarly positioned as the C-terminal amide, and the pyrazole ring aligns with P^34^ of PP (Figure 6C). Analysis of intermolecular distances revealed that the sidechains of seven key Y_4_R residues, including D6.59, N6.55, W107^ECL1^, H7.39, Q5.46, F7.35, and L6.51, are within 4.5 Å of Z9407529782 and are therefore favorably positioned to form interactions (***Mayol et al., 2019***), likely contributing to receptor activity. In contrast, the sidechains of six key Y_4_R residues, including E203, V3.36, T2.61, M7.43, F6.54, and I295, are outside this range, providing a structural rationale for the comparably lower activity of the small-molecule agonist compared to PP and related peptides (Figure 5B).

## Discussion

Obesity and related comorbidities represent one of the most challenging and rapidly growing global health issues (***GBD 2021 Adolescent BMI Collaborators, 2025***). As demonstrated by the clinical success of the GLP-1 agonist Semaglutide (***Salvador et al., 2025***) and the dual GLP-1/GIP agonist Tirzepatide (***Hamza et al., 2025***), targeting peptide-activated GPCRs has proven a highly effective treatment strategy (***Rubio-Herrera and Mera-Carreiro, 2025***). Notably, the Y_4_R represents an interesting alternative GPCR target that is critical in the regulation of satiety and food intake (***Sainsbury et al., 2010***). Thus, in the last decades, attempts have been undertaken to develop clinically applicable Y_4_R-selective peptides. Among full-length PP derivatives, including the dual Y_2_/Y_4_R agonist Obinepitide (**Schwartz TW**, WO/2005/089789. 2005) and PP 1420 (***Tan et al., 2012***), a PP analogue with an additional N-terminal glycine, a number of short peptides based on the C terminus of PP have been identified. These truncated peptides include the tandem peptides GR231118 (1229U91) (***Parker et al., 1998***), originally characterized as a Y_1_R antagonist, and BVD-74D (***Balasubramaniam et al., 2006***), which displays a low potency at Y_4_R. From these dimers, the monomeric tetrapeptide UR-AK32 (***Konieczny et al., 2020***) and hexapeptide UR-KK236 (***Kuhn et al., 2017***) were developed as partial Y_4_R agonists and further optimized via cyclization to yield the high-affinity peptides UR-AK95c and UR-AK86c (***Konieczny et al., 2021***). However, none of these peptides reached clinical approval, likely due to the low plasma stability of PP (***Adrian et al., 1978***) and its derivatives (***Bellmann-Sickert et al., 2011***; ***Konieczny et al., 2021***), highlighting the need for novel ligand discovery strategies.

In this study, PP and the cyclic hexapeptides UR-AK95c and UR-AK86c were used as molecular probes to investigate the molecular mechanisms underlying Y_4_R agonism (Figure 1A). Functional studies confirmed that the hexapeptides activate Y_4_R with a similar potency compared to the endogenous 36-mer PP (Figure 1B). This is in agreement with other cyclized, truncated GPCR peptide analogues, such as cyclic chemerin-9 (***Fischer et al., 2021***) or Octreotide (***Bauer et al., 1982***), which imitate the activity of their respective endogenous full-length ligands at CMKLR1 and somatostatin receptors, respectively. Notably, in our studies, a faster but less stable G-protein dissociation was observed upon hexapeptide-induced Y_4_R activation compared to full-length PP (Figure 1C). This behavior is consistent with their smaller size, which typically facilitates rapid association through faster diffusion and limited steric constraints (***Schreiber et al., 2009***), but also reduces residence time due to fewer stabilizing contacts (***Tresadern et al., 2011***).

In addition, it was shown that the improved binding selectivity of the hexapeptides for Y_4_R towards the other Y receptor subtypes also translates to functional selectivity (Figure 1D). The improved selectivity of UR-AK95c and UR-AK86c over PP is likely mediated by Y_4_R residues that selectively influence hexapeptide activity, including Y2.64, Q3.32, V200^ECL2^, E6.58, and ECL3 (Figure 3B, Figure 3-figure supplement 1). Sequence alignment (***Chenna et al., 2003***) across the four neuropeptide Y receptor subtypes reveals pronounced distinctions at these positions, with Y2.64 exchanged to serine in Y_5_R, E6.58 substituted to phenylalanine, valine, or threonine in Y_1_R, Y_2_R, and Y_5_R, respectively, and multiple variations in ECL3 (Figure 3 -figure supplement 3). These differences strongly suggest a prominent contribution of these receptor positions to the improved receptor subtype discrimination of the hexapeptides. UR-AK86c further displays modestly increased selectivity towards Y_2_R compared to UR-AK95c (Figure 1D). This is potentially caused by additional interactions between the hydroxyl group of Y_2_ and D2.68 (Figure 4D, E). Importantly, this residue is substituted by glycine in Y_2_R, underscoring position 2.68 as useful hotspot to finetune receptor subtype selectivity of NPY-related ligands (Figure 3-figure supplement 3).

The allosteric antagonist *(S)*-VU0637120 inhibits, whereas the positive allosteric modulator VU0506013 potentiates both the PP- and hexapeptide-induced receptor activation (Figure 1E). This conserved modulatory effect suggests an analogous positioning of the hexapeptides and the PP C terminus in the orthosteric Y_4_R pocket and is in line with the allosteric modulators’ proposed mechanism of action. *(S)*-VU0637120 partially overlaps with the orthosteric binding pocket and engages receptor residues of the key Y_4_R activation interface, e.g. W107^ECL1^ and F7.35, thereby interfering with binding of all three peptides (***Schüß et al., 2021***). In contrast, VU0506013 stabilizes the PP-Y^4^R complex by occupying a cavity between TM1, TM2, and TM7, where it forms hydrophobic contacts with P^34^ of PP (***Schüß et al., 2023***). Despite the replacement of this proline to leucine in the hexapeptides, the potentiation by VU0506013 is preserved, which suggests that L^4^ adopts a comparable orientation, enabling similar hydrophobic interactions with the PAM (***Schüß et al., 2026***).

Docking and mutagenesis studies determined 13 residues critical for stabilizing the active receptor conformation, which were defined as the key Y_4_R activation interface (Figure 4C). Strikingly, 11 of the 13 key residues are suggested to form contacts with the RXRY-NH_2_ motif, establishing it as main pharmacophore of the Y_4_R peptides (Figure 4C). T2.61 and H7.39 form polar contacts with the C-terminal amide of the peptides (Figure 4B, C). Notably, inactive Y_1_R structures revealed that the antagonists UR-MK299 and BMS-193885 omit contacts with T2.61 and H7.39, while blocking most of the interactions for the endogenous agonist NPY (***Yang et al., 2018***). This could indicate a prominent role of the residues T2.61 and H7.39 in triggering the activation of Y_1_R, but also of the phylogenetically closely related Y_4_R (***Larhammar and Salaneck, 2004***). Beyond these interactions, several additional key peptide-receptor contacts are consistently observed across all peptides: A hydrophobic patch (V3.36, L6.51, M7.43) interacts with the Y^36^/Y^6^ phenol ring, Q5.46 engages the Y^36^/Y^6^ hydroxyl group, an ionic/polar network (E203, N6.55, D6.59) coordinates R^35^/R^5^, and F6.54 and F7.35 form analogous contacts with R^33^/R^3^ (Figure 4B, C). Among these, T2.61, V/I3.36, E/D203, L6.51, F6.54, D6.59, F/Y7.35, H7.39, and M7.43 are conserved across all neuropeptide Y receptor subtypes, and the human Y_1_R and Y_4_R are even conserved at all identified positions, underscoring the critical role of these residues in mediating receptor activation (Figure 3-Figure supplement 3). Moreover, W107^ECL1^ was identified as functionally relevant for all three peptides. This tryptophan is part of the highly conserved WXXGXXXC motif found in class A GPCRs, which was previously associated with disulfide bridge formation (***Bhattacharyya et al., 2004***) or ligand guidance (***Jones et al., 2020***). Our findings expand this view by indicating that W107^ECL1^ contributes directly to ligand binding. In addition, the interaction between the α-helix of PP and the key Y_4_R residue I295^ECL3^ is likely mimicked by the macrocyclic structure of the hexapeptides (Figure 2-figure supplement 3). Consistently, the non-cyclic peptide RYRLRY-NH_2_ (***Konieczny et al., 2020***) displays reduced activity at Y_4_R compared to the cyclic peptide UR-AK86c (Figure 5), supporting a critical contribution of this macrocycle-ECL3 interaction to the stabilization of the active Y^4^R conformation.

Comparing different Y_4_R agonists further allowed us to distinguish residues required for receptor activation from those mediating ligand-specific interactions (Figure 4E). Y2.64, Q3.32, V200^ECL2^, and E6.58 were preferentially important for the hexapeptides, whereas D2.68, Y106^ECL1^, F184^ECL2^, and H188^ECL2^ contributed more prominently to PP signaling. L^4^ of the hexapeptides occupies a larger space around TM2 and TM3 compared to P^34^ of PP, rationalizing the higher sensitivity to the Y2.64A substitution (Figure 4E). Although Q3.32 forms polar contacts with the C-terminal amide in all ligands, its substitution selectively influences the hexapeptides, likely reflecting their smaller binding interface and reduced conformational flexibility due to cyclization (Figure 4B, E). E6.58 forms polar interactions with R^33^/R^3^ in all three peptides; however, only the hexapeptides are affected by its substitution to alanine (Figure 4E). This observation may be attributed to the cyclization in the hexapeptides, which constrains the conformation of R^3^, thereby reducing its flexibility and limiting its ability to form alternative interactions with the surrounding of E6.58 (Figure 4D). In addition, F184^ECL2^ and H188^ECL2^ are functionally relevant for PP but dispensable for the hexapeptides, whereas it is *vice versa* for V200^ECL2^ (Figure 4E). This distinct behavior likely reflects ligand flexibility: While L^24^ and I^28^ of PP are conformationally constrained in the α-helix, enforcing defined contacts with F184^ECL2^ and H188^ECL2^, W^2^/Y^2^ are more flexible, reducing their dependence on F184^ECL2^ and H188^ECL2^ and allowing for contacts with V200^ECL2^ (Figure 4D). In addition, the proposed interaction between T^32^/Y^2^ of PP/UR-AK86c and D2.68 is not mimicked by W^2^ of UR-AK95c (Figure 4D, E), highlighting D2.68 as a residue that serves as anchor for specific agonists, but does not trigger receptor activation.

Structure-activity studies of truncated UR-AK86c analogues revealed that shortening of the R^5^ sidechain reduces peptide activity, which could be due to a destabilization of contacts with E203^ECL2^, N6.55, and D6.59, highlighting the prominent role of interactions with this ionic/polar network in Y_4_R activation (Figure 5). Similarly, deletion of the cyclic linker diminished peptide activity, likely reflecting reduced affinity due to the loss of conformational constraint and the resulting increase of entropic penalty of receptor binding (***Gaucher et al., 2022***). However, the loss of function of RYRLRY-NH₂ relative to UR-AK95c is observed exclusively at Y_4_R, but not at the remaining receptor subtypes, suggesting that the cyclic linker also plays a role in interactions with specific residues unique for Y_4_R. Combined docking and receptor loop chimera experiments indicate that these interactions likely involve ECL3, highlighting ECL3 as receptor segment that can be exploited to achieve Y_4_R-selectivity (Figure 2-figure supplement 3, Figure 3-figure supplement 1). Intriguingly, the tetrapeptide RLRY-NH_2_ still activates Y_4_R with nanomolar potency (Figure 5). We suggest that the selective responsiveness of Y_4_R to very small peptides is due to a low energy barrier for receptor activation, which is also evident in the higher constitutive activity of Y_4_R compared to the other neuropeptide Y receptor subtypes (***Chen et al., 2000***). This characteristic makes Y_4_R a promising target for small-molecule agonists.

Guided ULLS, informed by the key Y_4_R interface, identified three selective non-peptide small-molecule agonists for Y_4_R (Figure 6A). Redocking of the most potent compound Z9407529782 revealed proximity to several residues critical for receptor activation, indicating a partial imitation of the native peptide interaction network. However, key contacts are not fulfilled, e.g. with T2.61, which caused a substantial 100-1000-fold loss in potency for PP and the hexapeptides. Structural modification of the hit to address this residue could therefore improve potency and provides a clear direction for structure-guided optimization. Remarkably, the development of Danuglipron (GLP1R) (***Griffith et al., 2022***) and PCO371 (PTH1R) (***Tamura et al., 2016***) illustrate that potent agonism at peptide-binding GPCRs, which usually feature a broad ligand-receptor interface (***Wu et al., 2017***), can be achieved through non-peptide small-molecules.

In summary, this study combined experimental and computational approaches to elucidate the pharmacophoric key interface of Y_4_R. We believe that these insights provide a valuable basis for the rational optimization of the hit identified in this study into potent and selective small-molecule Y_4_R agonists, as this GPCR appears to be activatable by remarkably short peptides. In the future, such compounds could expand the molecular toolbox for Y_4_R to support the investigation of metabolic disorders and the development of novel therapeutics targeting obesity.

## Material and Methods

### Generation of plasmids

The cDNA of wild-type Y_4_R and Y_1_R was cloned into a pEYFP_N1 vector (Clontech, Heidelberg, Germany), resulting in receptor constructs C-terminally fused to enhanced yellow fluorescent protein (eYFP), as previously described (***Dinger et al., 2003***). For untagged Y_4_R, UAA stop codon was introduced upstream of eYFP in Y_4_R_eYFP_N1 plasmid by Q5^®^ site-directed mutagenesis PCR. The chimeric G protein Δ6Gα_qi4myr_ cDNA was generously provided by E. Kostenis (Rhenish Friedrich Wilhelm University of Bonn, Germany) (Kostenis, 2001). The G_i3_ protein-based, tricistronic activity sensor (G_i3_-CASE) plasmid was developed by H. Schihada (Philipps University of Marburg, Germany) (***Schihada et al., 2021***) and generously provided by G. Schulte (Karolinska Institute, Solna, Sweden). The sequences of all constructs were validated by Sanger sequencing (Microsynth Seqlab, Göttingen, Germany).

### Generation of Y_4_R residue variants and Y_4_/Y_1_R chimeras

Single- and multiple-residue receptor variants were generated by introducing amino acid substitutions into wild-type or mutant Y_4_R_eYFP_N1 using the Q5^®^ mutagenesis kit (New England Biolabs, Ipswich, MA, USA), as described previously (***Schermeng et al., 2025***). Primers were designed using NEBaseChanger. To phosphorylate and ligate the linear PCR products and digest the parental DNA, KLD Enzyme Mix in KLD Reaction buffer (New England Biolabs, Ipswich, MA, USA) was used according to the manufacturer’s protocol. Y_4_/Y_1_R chimeras were produced by employing overlap extension PCR (***Ho et al., 1989***). The detailed process was previously described (***Schüß et al., 2021***). Briefly, the N-terminal cDNA of the donor receptor subtype was obtained in a PCR using a sense primer with a specific restriction site and an antisense primer with an overhang complementary to the cDNA of the second receptor subtype. In a second PCR, the C-terminal cDNA of the second receptor subtype was amplified, using a sense primer with an overhang complementary to the N-terminal cDNA of the donor receptor and an antisense primer with a restriction site. Both PCR products were separated by agarose gel electrophoresis and purified using Promega Wizard® SV Gel and PCR Clean-Up System (Promega, Madison, WI, USA). The last PCR combined the fragments to a chimeric cDNA. The final product was digested with respective restriction enzymes, purified as described above, and ligated into a pEYFP_N1 vector using T4 DNA ligase. PCR products from site-directed mutagenesis PCR and overlap-extension PCR were amplified in *Escherichia coli* DH5α and purified using PureYield^TM^ Plasmid Midiprep System (Promega, Madison, WI, USA), as described previously (***Schüß et al., 2021***). The concentration of plasmid solutions was determined using a multimode microplate reader Infinite^®^ M200 (Tecan, Männedorf, Switzerland). The sequences of all constructs were validated by Sanger sequencing (Microsynth Seqlab, Göttingen, Germany).

### Cell culture

COS-7 cells were purchased from the American Type Culture Collection (ATCC) and maintained in Dulbecco’s modified Eagle’s medium (DMEM, Biowest, Nuaillé, France), supplemented with 10 % sterile filtered fetal bovine serum (FBS, Sigma-Aldrich, St. Louis, MO, USA). As described in detail before, stably co-transfected COS-7_hY_1/2/4/5_R_ΔGα_qi4myr_ cells were generated using linearized pVitro2-MCS vectors (Invivogen, San Diego, CA, USA) (***Mäde et al., 2014***). The first vector carried Y_4_R_eYFP cDNA and a hygromycin resistance gene, while the second vector contained Δ6Gα_qi4myr_ cDNA and a G418 resistance gene. They were maintained under selective conditions in DMEM, supplemented with 10 % FBS, 133 µg/mL hygromycin, and 1.5 mg/mL G418 sulfate (Invivogen, San Diego, CA, USA). HEK293 cells were purchased from Deutsche Sammlung von Mikroorganismen und Zellkulturen (DSMZ, Braunschweig, Germany) and maintained in DMEM/Ham’s F-12 (1:1 (v/v), Biowest, Nuaillé, France), supplemented with 15 % FBS. All cell lines were cultured under a humidified atmosphere at 37 °C and 5 % CO_2_ until confluency, and passaged twice a week using trypsin/ethylenediaminetetraacetic acid (EDTA, Biowest, Nuaillé, France) for detachment.

### IP-one accumulation assay

For the functional characterization of the peptides and compounds at neuropeptide Y receptor subtypes, IP-one accumulation assay was conducted in stably co-transfected COS-7_Y_1/2/4/5_R_ΔGα_qi4myr_ cells, as described recently (***Schüß et al., 2021***). COS-7_Y_1_R_ΔGα_qi4myr_ cells and COS-7_Y_5_R_ΔGα_qi4myr_ cells were seeded with a density of 8,000 cells, whereas COS-7_Y_2_R_ΔGα_qi4myr_ cells and COS-7_Y_4_R_ΔGα_qi4myr_ were seeded with a density of 4,000 cells and incubated under a humidified atmosphere at 37 °C and 5 % CO_2_ for 24 h. For mutagenesis studies, the assay was performed with COS-7 cells transiently transfected with wild-type or mutant receptor-eYFP constructs and chimeric G protein in a 3:1 ratio. The protocol was published previously and adapted (***Tang et al., 2022***). Cells were transfected using Metafectene^®^ Pro transfection reagent (Biontex, Munich, Germany) according to the manufacturer’s protocol and seeded with a density of 8,000 cells in a 384-well plate (Greiner, Kremsmünster, Austria). After incubation of 24 h under a humidified atmosphere at 37 °C and 5 % CO_2_, the medium was removed and peptide-induced IP-one accumulation was assessed using IP-one Gq Detection Kit (Revvity, Waltham, MA, USA) according to the manufacturer’s protocol. Briefly, cells were incubated for 45 min with the peptide, after which fluorescently labelled IP_1_-d2 (HTRF acceptor) in lysis buffer was added, followed by the addition of IP_1_ Tb cryptate antibody (HTRF donor) in lysis buffer. Subsequently, plates were shaken at room temperature for 1 h, and fluorescence emission was measured at 620 nm (HTRF donor) and 665 nm (HTRF acceptor) using a Spark® multimode microplate reader (Tecan, Männedorf, Switzerland). The HTRF ratio for each well was calculated by dividing the fluorescence intensity at 665 nm by the fluorescence intensity at 620 nm and multiplying the quotient by a factor of 10,000. The HTRF ratio was further analyzed using GraphPad Prism 10.5.0 (GraphPad Software, San Diego, CA, USA).

### BRET-based G_i3_ protein assay

For the functional characterization of the peptides at Y_4_R, a bioluminescence resonance energy transfer (BRET)-based G_i3_ protein assay was performed. The assay was conducted in COS-7 cells transiently co-transfected with untagged Y_4_R and G_i3_-CASE in a 1:1 ratio. G^i3^-CASE plasmid encodes all three subunits of the G^i3^ protein, with Gα fused to NanoLuc (Nluc) nanoluciferase and Gγ fused to fluorescent protein cpVenus (***Schihada et al., 2021***). COS-7 cells were co-transfected with Metafectene^®^ Pro transfection reagent (Biontex, Munich, Germany) according to the manufacturer’s protocol. Subsequently, cells were resuspended in DMEM/Ham’s F-12 (phenol red-free, Gibco Thermo Fisher Scientific, Waltham, MA, USA), supplemented with 10 % FBS. A total of 100 µL (50,000 cells) were seeded into each well of a white, solid 96-well plate (Greiner, Kremsmünster, Austria). After at least 24 h incubation under a humidified atmosphere at 37 °C and 5 % CO_2_, medium was aspirated from each well and replaced by 100 µL BRET buffer (HBSS, 4-(2-hydroxyethyl)-1-piperazineethanesulfonic acid (HEPES, Sigma-Aldrich, St. Louis, USA), pH 7.2 at 37°C, sterile filtered). Fifty µL coelenterazine h solution (NanoLight Technology, Pinetop, AZ, USA) was added with a final concentration of 4.2 µM. Cells were incubated for 5 min at 37 °C. For the generation of concentration-response curves, cells were stimulated for 15 min at 37 °C with 50 µL per well of a dilution series of the peptides. BRET buffer without peptide served as baseline control. For the kinetic studies of the peptide-induced dissociation of the G protein sensor, Nluc luminescence and cpVenus fluorescence was measured for 5 min after addition of coelenterazine h. Subsequently, two concentrations of peptide and buffer as baseline control were added. The BRET measurement was continued for 30 min. Nluc luminescence (400 – 470 nm) and cpVenus fluorescence (535 – 650 nm) was determined using multimode microplate reader Spark^®^ (Tecan, Männedorf, Switzerland). The BRET ratio of each well was calculated by dividing cpVenus emission intensity by Nluc emission intensity. To obtain the netBRET value, the average BRET ratio value of the baseline control was substracted. The calculated netBRET values were analyzed using GraphPad Prism 10.5.0.

### Ca^2+^-flux assay

The impact of the positive allosteric Y_4_R modulator VU0506013 and the allosteric Y_4_R antagonist (*S*)-VU0637120 on the activity of PP, UR-AK95c, and UR-AK86c was investigated using Ca^2+^-flux assay in stably co-transfected COS-7_Y_4_R_ΔGα_qi4myr_ cells, as previously described (***Schüß et al., 2026***). Cells were seeded with a density of 30,000 cells per well in black clear bottom 96-well plates (Greiner, Kremsmünster, Austria). After incubation of 24 h under a humidified atmosphere at 37 °C and 5 % CO_2_, Ca^2+^-flux assay was conducted, as previously described (***Schubert et al., 2017***). In short, COS-7_Y_4_R_ΔGα_qi4myr_ cells were labeled with 2.4 µM Fluo-2 AM calcium dye (Abcam, Cambridge, United Kingdom) in assay buffer (20 mM HEPES, 2.5 mM probenecid (Sigma-Aldrich, St. Louis, MO, USA) in HBSS, pH 7.4) for 60 min. After replacement of labeling solution by assay buffer, basal Ca^2+^ levels (excitation 485nm, emission 525 nm) were detected for 20 s using a Flex Station III device (Molecular Devices, San Jose, CA, USA) Subsequently, DMSO (Sigma-Aldrich, St. Louis, MO, USA) or compound solution was added for 60 s, followed by the addition of peptide solution and measured for further 70 s. The raw data was processed to x-fold over basal and normalized to the maximum and minimum values of the DMSO control. Additional analysis steps were performed using GraphPad Prism 10.5.0.

### Live cell fluorescence microscopy

To investigate the expression and subcellular localization of wild-type and mutant Y_4_R variants, HEK293 cells expressing the receptor-eYFP constructs were examined in fluorescence microscopy, as previously reported (***Rathmann et al., 2013***). Cell nuclei were stained with HOECHST33342 (Sigma-Aldrich, St. Louis, MO, USA). Cells were imaged using an ApoTome Imaging System with an Axio Observer.Z1 microscope (Carl Zeiss, Oberkochen, Germany) equipped with 63x/1.40 oil objective and two filters (YFP: filter set 46; 4′,6-diamidino-2-phenylindole (DAPI): filter set 49). Microscopy images were processed using Zen3.6 (blue edition, Carl Zeiss, Oberkochen, Germany).

### Peptide synthesis

Human pancreatic polypeptide (hPP, sequence: APLEPVYPGDNATPEQMAQYAADLRRYINMLTRPRY-NH_2_), porcine neuropeptide Y (pNPY, YPSKPDNPGEDAPAEDLARYYSALRHYINLITRQRY-NH_2_), the hexapeptide RYRLRY-NH_2_, and the tetrapeptide RLRY-NH_2_ (UR-AK32) were synthesized by automated solid-phase peptide synthesis in a multiple peptide synthesis robot (Syro I, MultiSyntech, Witten, Germany) using a 9-fluorenylmethyloxycarbonyl/tert-butyl (Fmoc/tBu) strategy and purified using preparative reversed-phase high-performance liquid chromatography (RP-HPLC) on an Aeris PEPTIDE XB-C18 column (100 Å, 5 μm, 250 × 2.1mm^2^, Phenomenex, Torrance, CA, USA), as described earlier (***Pedragosa-Badia et al., 2014***). UR-AK95c, UR-AK86c, and [Agp^5^]-UR-AK86c were generously provided by Prof. Dr. Max Keller (University of Regensburg, Germany). The detailed synthesis protocol of UR-AK95c and UR-AK86c was previously published (***Konieczny et al., 2021***). The detailed synthesis protocol of [Agp^5^]-UR-AK86c is included in the Supplementary File.

Purity ≥ 95 % was ascertained using two different analytical RP-HPLC systems (Jupiter Proteo 90 Å C12 and Aeris Peptide 100 Å XB-C18, Phenomenex, Torrence, USA) with linear gradients of 5 – 55 %, 10 – 60 %, or 20 – 70 % eluent B (0.1 % TFA in ACN) in eluent A (0.08 % TFA in H_2_O) in 40 min, respectively. Matrix assisted laser desorption ionization - time of flight mass spectrometry (MALDI-ToF, UltrafleXtreme, MALDI-ToF/ToF, Bruker Daltonics, Billerica, MA, USA) was used to confirm the identity of the peptides (Figure 5-figure supplement 1).

### Compound synthesis

The positive allosteric Y_4_R modulator VU0506013 was synthesized and purified, as described previously (***Schüß et al., 2023***). The allosteric Y_4_R antagonist (*S*)-VU0637120 was synthesized and purified, as recently described (***Schüß et al., 2021***). The identity of both compounds was determined using ^1^H-NMR. The purity of ≥ 95 % was ascertained using RP-HPLC. (Figure 5-figure supplement 1) Selected compounds from the ULLS were ordered and synthesized by Enamine LTD using their in-house pipelines (***Grygorenko et al., 2020***). Compounds were produced in three successive batches comprising 14, 30, and 9 molecules, respectively, yielding a total of 53 compounds. A complete SMILES list of the ordered and tested compound is included in the Figure 6 -table supplement 1. All ordered compounds could be successfully synthesized. Stock solutions were prepared by dissolving approximately 5 mg of each compound in dimethyl sulfoxide (DMSO). Compound identity and purity were confirmed by liquid chromatography–mass spectrometry (LC–MS) analysis performed by the supplier, with all compounds meeting a purity threshold of ≥90 %.

### Computational methods

#### Y_4_R and PP modelling

The UR-AK95c and UR-AK86c structures were generated using Rosetta3.14 with custom cross-linking methods (***Leman et al., 2020***). The cryo-EM structure of Y_4_R with bound PP was obtained from the Protein Data Bank (PDB: 7X9C) (***Tang et al., 2022***) and used as the basis for modelling. Water, ions, lipids as well as the bound G-protein were stripped and the structure energetically minimized using the Rosetta relaxation protocol (***Nivón et al., 2013***) to generate 500 structures. The best-scoring structure by total energy was selected and the bound PP was reduced to the last six amino acids to fit the length of the cyclic peptide. This minimized structure was used as a reference point for modelling of the hexapeptides. To address the intrinsic flexibility of the receptor loops, the protein structure was again minimized with restricted helix movement as well as restricted movement for the intracellular loops to generate an ensemble of flexible extracellular loops. 1,000 structures were sampled and the 400 top-scoring structures were taken, adopting a threshold of -790 Rosetta Energy Units (REU) to remove models that did not converge in energy. These structures were then clustered using a k-means algorithm with total energy and RMSD for the best-scoring structure as the clustering parameters. Six clusters were generated, and their representatives were used in the computational screening section of the study.

##### Generation of cyclic peptides

The cyclic peptides were generated using Rosetta’s Crosslink mover (***Fleishman et al., 2011***). The initial structures of the linear peptides were based on the PP structure, which was modified by mutating position L^1^ to R^1^, P^4^ to L^4^ to get the linear precursor of UR-AK95c. In addition, position W^2^ was mutated to Y^2^ to create an initial structure of the linear precursor of UR-AK86c. The cyclization was implemented computationally in three steps: adding a patch to R^1^, modifying the R^3^ side chain (ARC residue), and adding a succinyl linker between residues X and Y (SCN). Parameter files for the linker and ARC residue were generated using a combination of RDKit (***Landrum, 2013***), the fake_rotlib.py script (***Bell et al., 2025***) and the molfile_to_params_polymer.py script as described in the Protocol Capture in the Supplementary File. For ARC, 1,000,000 conformations were generated, of which 100 conformations were included in RDKit based on MMFF scoring (***Wang et al., 2020***). For the linker, we used BCL and the molfile_to_params.py script. From 500 conformations, the 50 best scoring ones were selected based on BCL (***Brown et al., 2022***) and used to generate a params file.

### Ensemble docking of cyclic peptides

The docking of UR-AK95c and UR-AK86c into Y_4_R was carried out with the FlexPepDock algorithm using the ref2015_cart_cst scoring function (***Raveh et al., 2010***). The initial peptide arrangement was based on the PP backbone in the experimentally determined structure and prepared with Rosetta scripts (***Fleishman et al., 2011***). As a pre-selection step, we conducted 1,000 docking runs for each of the selected cluster representatives of the Y_4_R structure for each peptide. Based on the initial results, two structures were removed due to their poor performance and clashing of residues, resulting in unfavourable scores. In the following main docking, the remaining four structures were sampled 10,000 times each. Each generated structure was rescored with ref2015 to assess the quality of the bound peptides and remove those with poor scoring decoys.

### Selection of the best structures

To select the best-fitting models, a combined approach involving computational scoring and the integration of experimental data filter was applied. Here, the best-scoring models in both total energy and interface scores were selected and evaluated using the Rosetta energy breakdown algorithm (energetic hotspot analysis or contact map). Peptide-receptor contacts with energies below the threshold of -0.1 REU were defined as interactions. Models were ranked based on the number of experimentally observed interactions that could be found in the computational models. A fraction of these interactions was then used to select the models. The selected models were visualized using PyMOL (ver. 2.5.4) to verify their correlation with the experimental data and hotspot analysis.

### Ultra-large high-throughput screening

#### Preparation of compound database

The combinatorial library specification for the Enamine REAL chemical space was obtained directly from Enamine Ltd. under an academic non-disclosure agreement. Every compound in the REAL database can realistically be synthesized within a few weeks using established building blocks and validated reaction pathways. The library covers an exceptionally broad chemical space with diverse scaffolds, functional groups, and molecular frameworks, supporting extensive exploration during virtual screening. This database consisted of lists of reactions (building rules) and associated reactants (building blocks/substrates). When fully enumerated these combinations add up to 35,519,219,406. Reactions with two and three substrates were defined in SMARTS format (building rules) and substrates with their defined usage in the building rules were provided in SMILES format. All SMARTS and SMILES were combined into two tab-separated text files and provided as input for REvoLd (***Eisenhuth et al., 2025***). Virtual compound enumeration was performed using RDKit (version 2022.09.1) (***Landrum, 2013***), which combined substrates into fully specified molecules according to the defined building.

For all redocking steps, selected compounds and their corresponding two-dimensional representations were prepared following established BioChemical Library (BCL; version 4.3.0) (***Mendenhall et al., 2021***) and RosettaLigand protocols (***DeLuca et al., 2015***; ***Lemmon and Meiler, 2012***; ***Meiler and Baker, 2006***), using Rosetta (version 2022.45–3.13) and the REvoLd-adapted Rosetta release (version 2021.40).

### ULLS and filtering

To identify putative small-molecule Y_4_R agonists, the evolutionary docking algorithm REvoLd was applied as previously described (***Eisenhuth et al., 2025***), suing the standard REvoLd protocol without modification. REvoLd requires as input a receptor structure, a predefined binding-site region, and a combinatorial chemical library. In this study, the Enamine REAL chemical space served as the ligand source and the orthosteric binding site of the bound PP.

Following selection and validation of the Y_4_R receptor models, REvoLd was used to iteratively sample and optimize small-molecules from the Enamine REAL database against each receptor structure. For each receptor model, approximately 280,000 candidate compounds were generated and docked.

The following filtering workflow was applied as described before (***Liessmann et al., 2025***). In short: Initial filtering was applied to retain compounds with drug-like physicochemical properties (derived from Lipinski’s Rules of Five) (***Lipinski et al., 1997***), including molecular weight between 150 and 500 Da, calculated logP between −1 and 5, no more than 10 rotatable bonds, and no more than 5 hydrogen bond donors and 12 hydrogen bond acceptors. All physicochemical properties were computed using RDKit (version 2022.09.1). In addition, compounds containing Pan Assay Interference Compounds (PAINS) substructures were excluded using the RDKit PAINS filter catalog The best scoring 57,000 compounds from the evolutionary sampling stage were subjected to redocking. Here, BCL (version 4.3.0) was applied to generate multiple low-energy conformers per ligand, which were subsequently redocked into the Y_4_R binding site using RosettaLigand to refine binding poses and rescoring. Compounds that consistently ranked among the top candidates after redocking were retained. Here, a score-cutoff of -3.5 lid_root2 was applied (the interface energy normalized for molecule size by dividing the score with the square root of the number of non-hydrogen atoms), resulting in a subset of 9,512 ligands with highly favorable predicted interaction energies.

To minimize off-target activity within the neuropeptide Y receptor family, compounds were counter-screened *in silico* against the closely related Y_1_R and Y_5_R. Only ligands predicted to selectively bind Y_4_R, while lacking favorable binding to Y_1_R or Y_5_R, were retained, reducing the compound set to 574 Y_4_R-selective candidates.

To ensure chemical diversity, k-means clustering (100 clusters) was performed based on ligand structural features. Representative compounds from each cluster were selected using an RMSD versus binding energy analysis, which assessed the consistency of binding poses relative to the best-scoring conformation. Compounds displaying both favorable binding energies and stable binding geometries were prioritized. Further details on the filtering procedure were previously described (***Liessmann et al., 2025***).

Final compound selection was based on (i) the quality of the predicted binding mode within the Y4R binding pocket, including deep pocket engagement and the presence of at least one potential hydrogen bond, and (ii) proximity to key Y4R trigger residues identified in the mutagenesis studies.

#### Redocking of Z9407529782

After the activity of Z9407529782 had been confirmed, the compound was redocked to elucidate its potential binding modes. A total of 200 ligand conformers were generated with OMEGA (***Hawkins et al., 2010***) and subsequently docked with RosettaLigand (***Lemmon and Meiler, 2012***), yielding 1,000 docking poses. The top 30 poses were visually inspected and filtered by contact pattern with Y_4_R residues known to contribute to receptor activation. From these, top 5 scoring poses were selected and analyzed for ligand strain using the TorsionAnalyzer (***Penner et al., 2022***). Because the sulfonamide bond adopted angles that deviated slightly from experimentally observed values reported in the Cambridge Structural Database (CSD), and to account for potential induced-fit effects involving backbone and side-chain rearrangements, the five selected poses were further minimized using MOE (***Chemical Computing Group ULC, 2026***). This procedure resulted in the potential binding mode shown in Figure 6.

## Supplements

**Figure 1 -table supplement 1:**
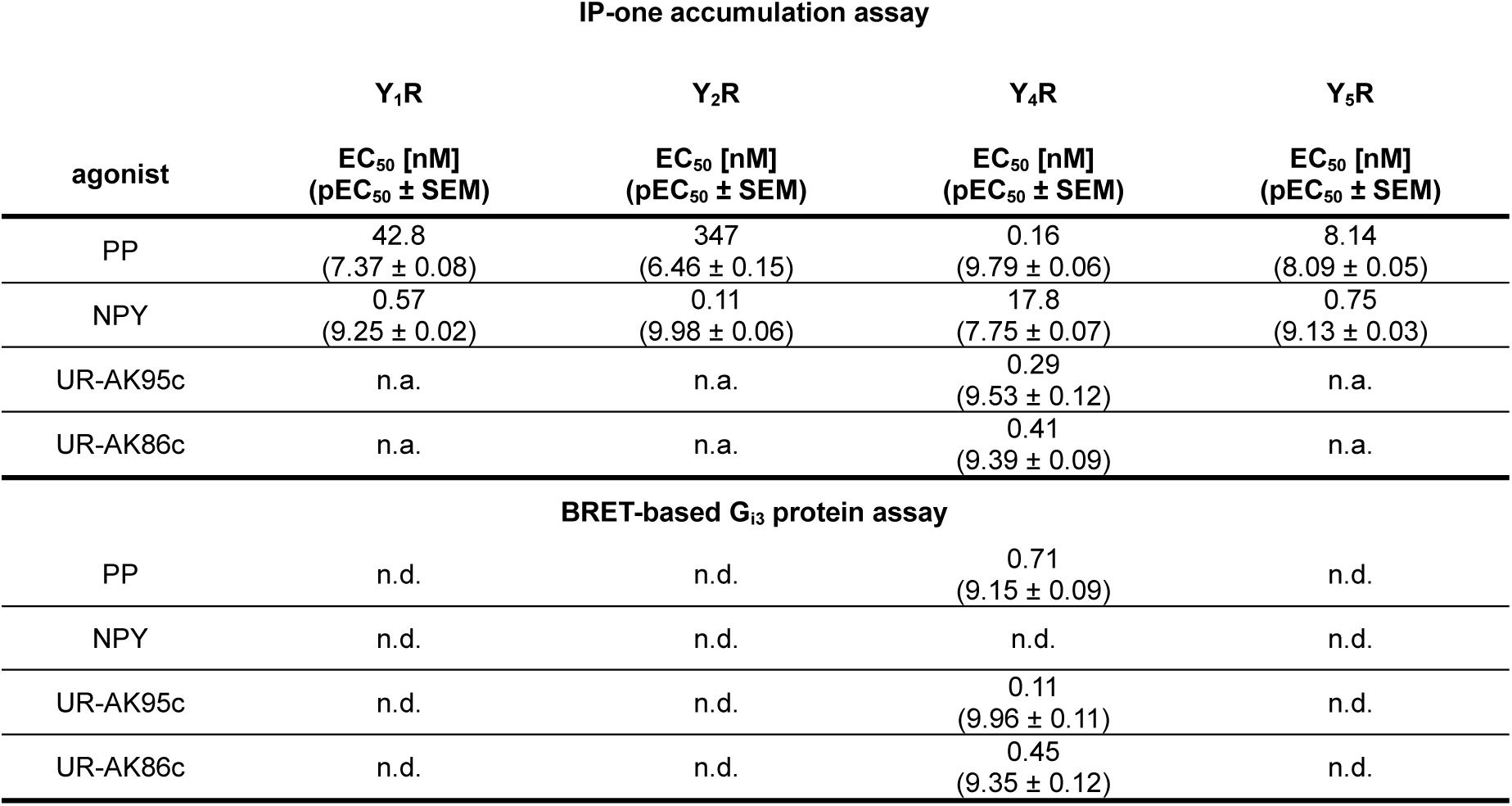
Functional data of tested peptides obtained by IP-one accumulation and BRET-based Gi3 protein assays. IP-one accumulation assay was performed in COS-7 cells stably expressing the Y1/2/4/5R-eYFP construct and the chimeric G protein Δ6Gαqi4myr. BRET-based Gi3 protein assay was performed in COS-7 cells transiently co-transfected with untagged Y4R and Gi3-CASE construct. EC50 and pEC50 values were calculated from a minimum of three independent experiments. n.a., not applicable, n.d., not determined.

**Figure 1 -table supplement 2:**
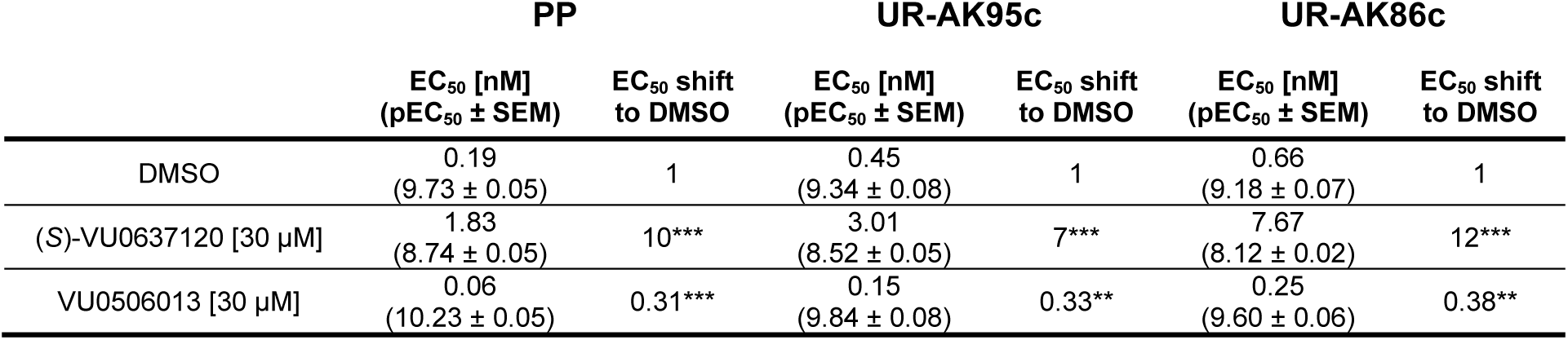
Functional characterization of PP, UR-AK95c, and UR-AK86c at Y4R in the presence of DMSO, 30 µM (*S*)-VU0637120, and 30 µM VU0506013. EC50 values were determined using Ca^2+^-flux assay in COS-7 cells stably expressing the Y4R-eYFP construct and the chimeric G protein Δ6Gαqi4myr. The EC50 shift represents the ratio of the EC50 value of peptide in the presence of an allosteric modulator to the EC50 value of the same peptide in the presence of DMSO. Each condition was tested in at least three independent experiments. Statistical analysis was conducted using one-way ANOVA followed by Dunnett’s post-test. n.a., not applicable, *, P < 0.033, **, P < 0.002, ***, P < 0.001.

**Figure 2 -figure supplement 1:**
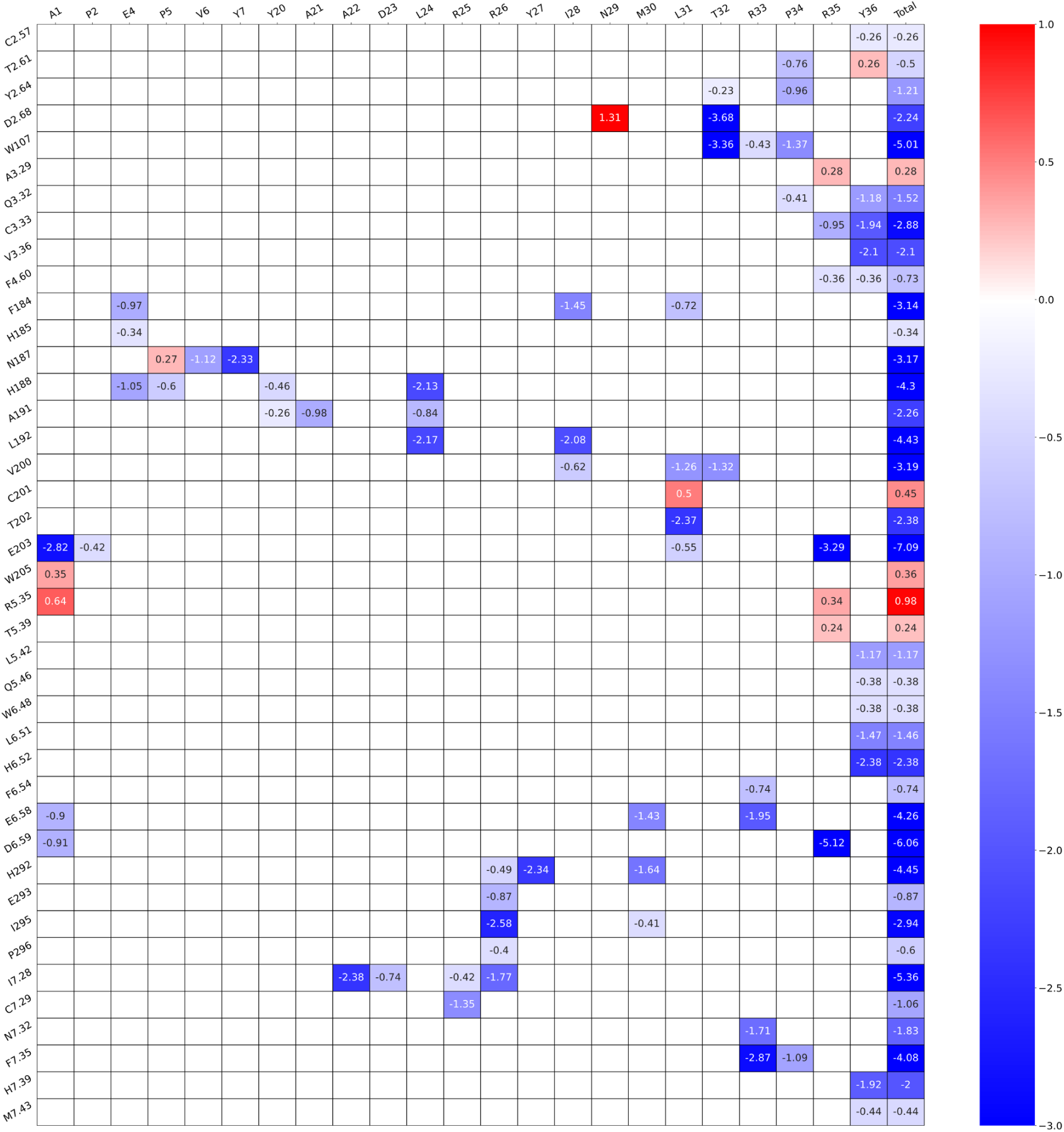
Contact map of per-residue energy breakdown of the best-fitting model of PP at Y4R. Average pairwise energy scores are depicted in Rosetta Energy Units (REU). The residues are named according to Ballesteros-Weinstein nomenclature (***Ballesteros and Weinstein, 1995***).

**Figure 2 -figure supplement 2:**
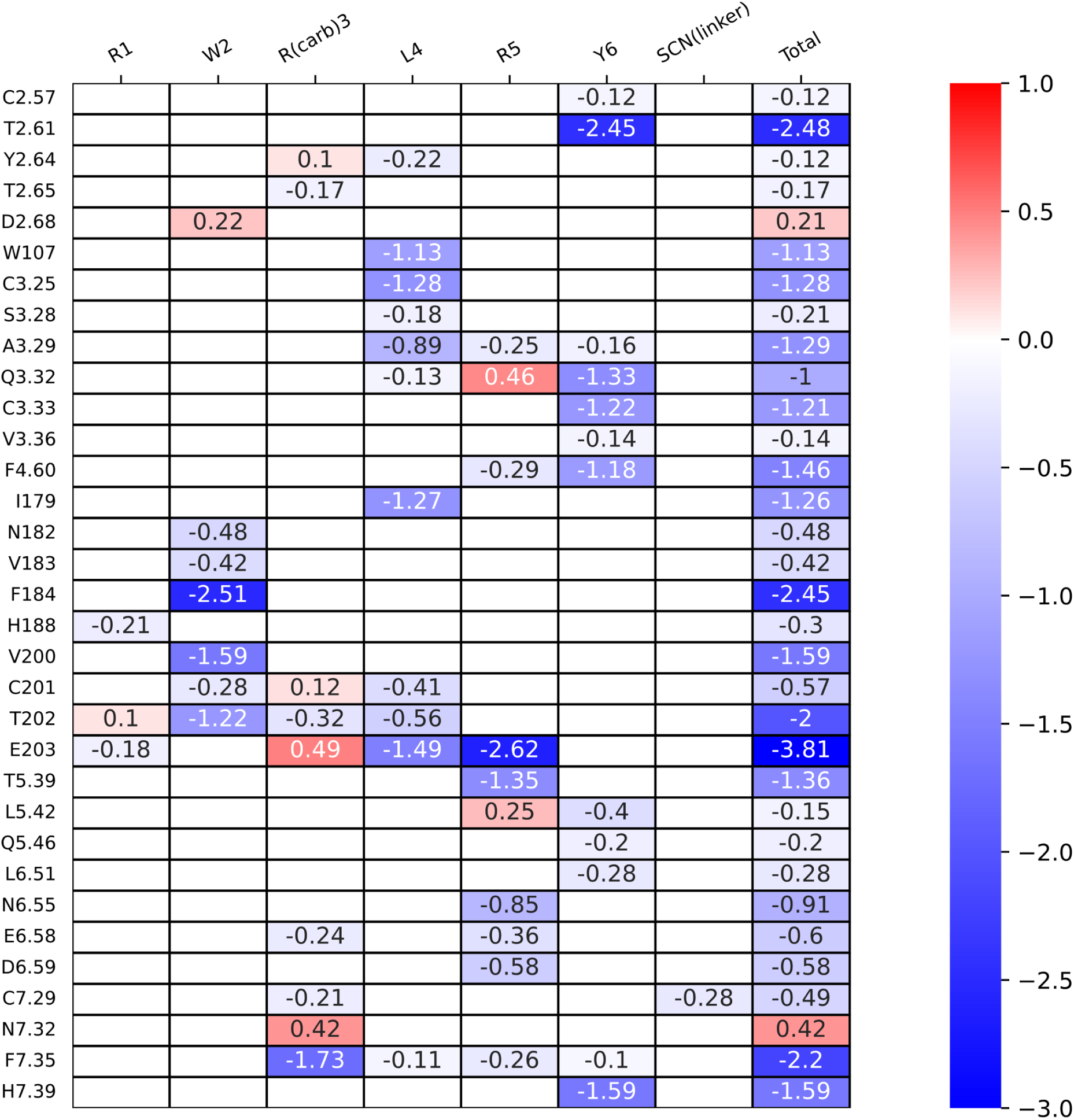
Contact map of per-residue energy breakdown of the best-fitting model of UR-AK95c at Y4R. Average pairwise energy scores are depicted in Rosetta Energy Units (REU). The residues are named according to Ballesteros-Weinstein nomenclature (***Ballesteros and Weinstein, 1995***).

**Figure 2 -figure supplement 3:**
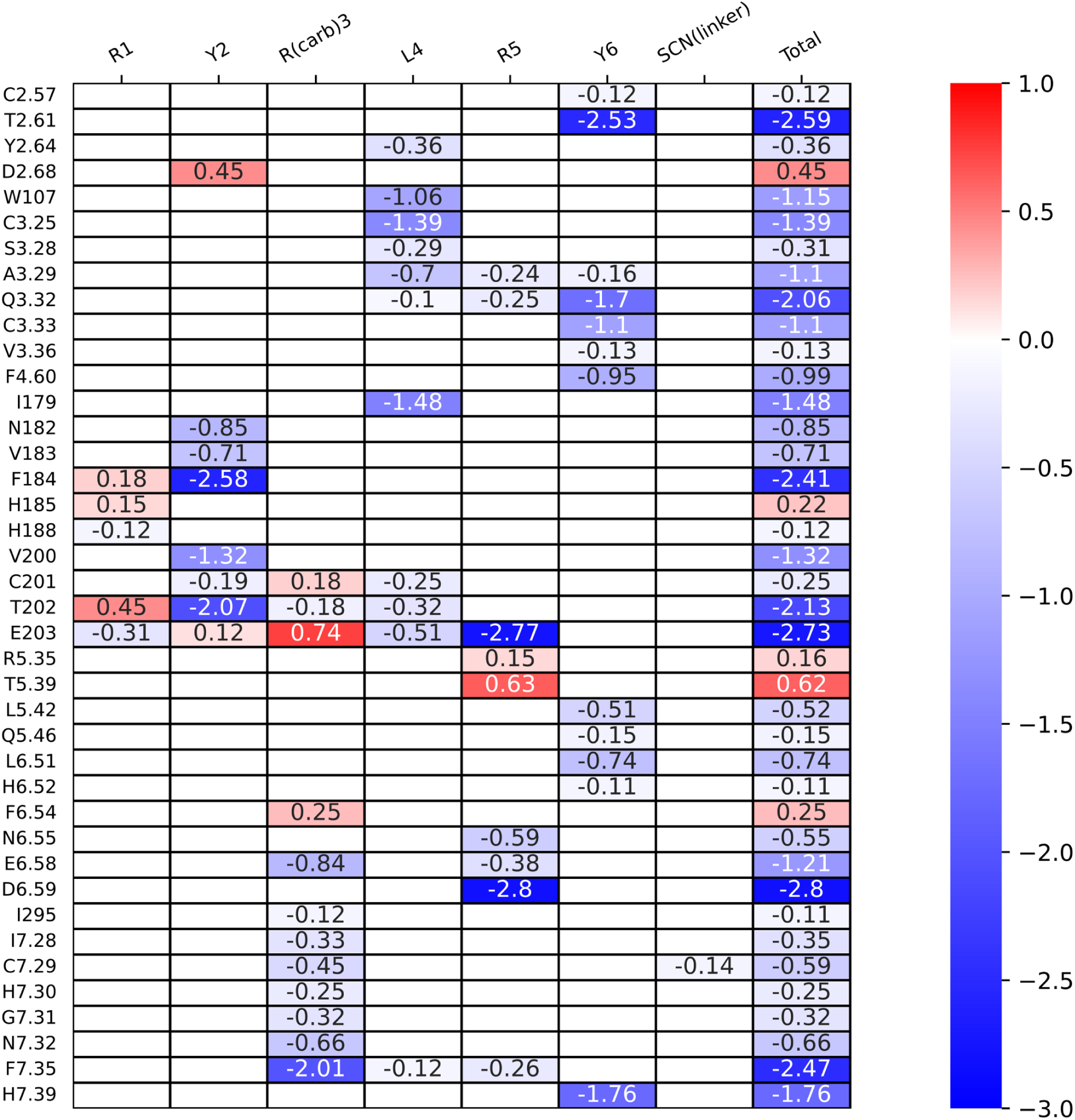
Contact map of per-residue energy breakdown of the best-fitting model of UR-AK86c at Y4R. Average pairwise energy scores are depicted in Rosetta Energy Units (REU). The residues are named according to Ballesteros-Weinstein nomenclature (***Ballesteros and Weinstein, 1995***).

**Figure 3 -table supplement 1:**
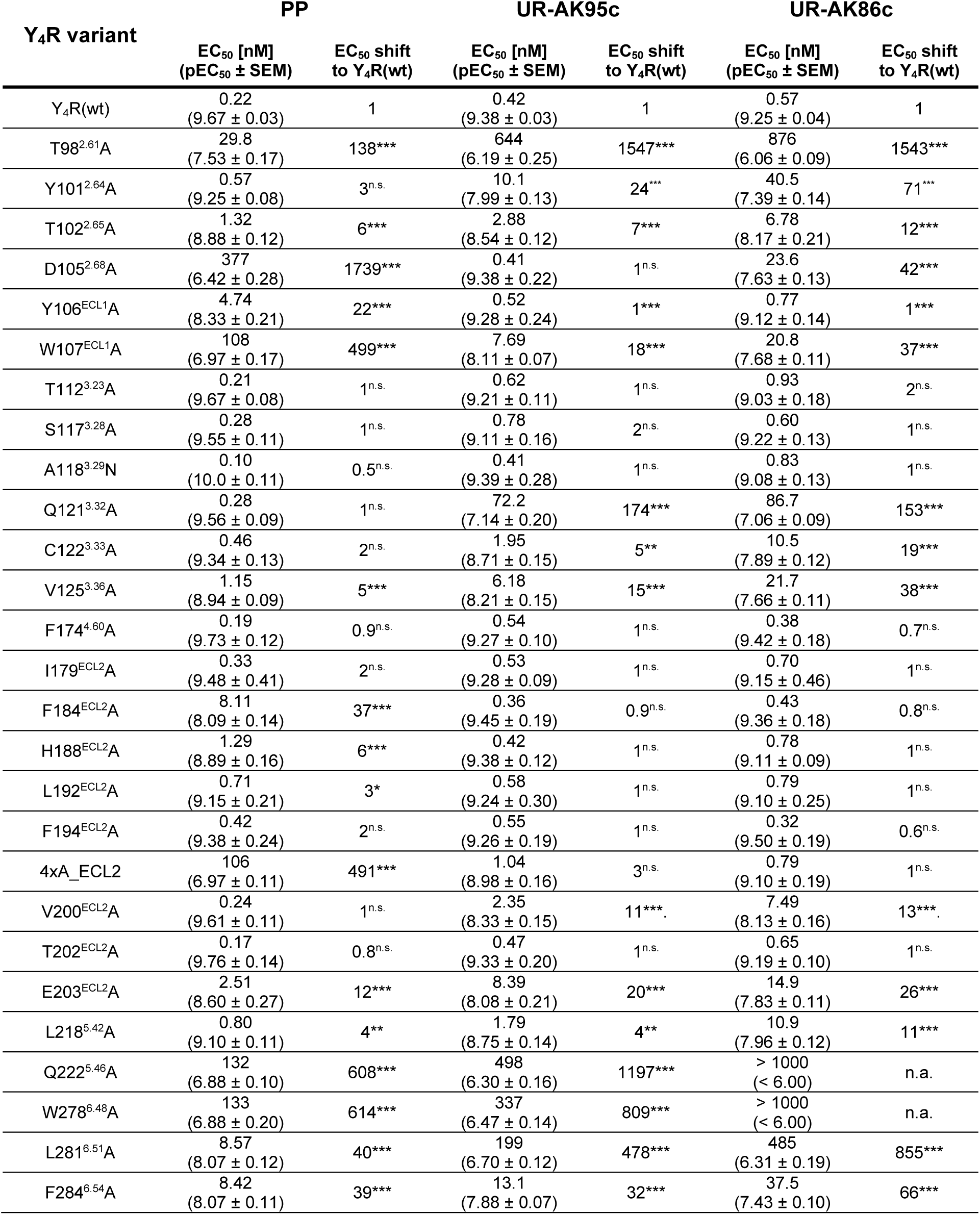
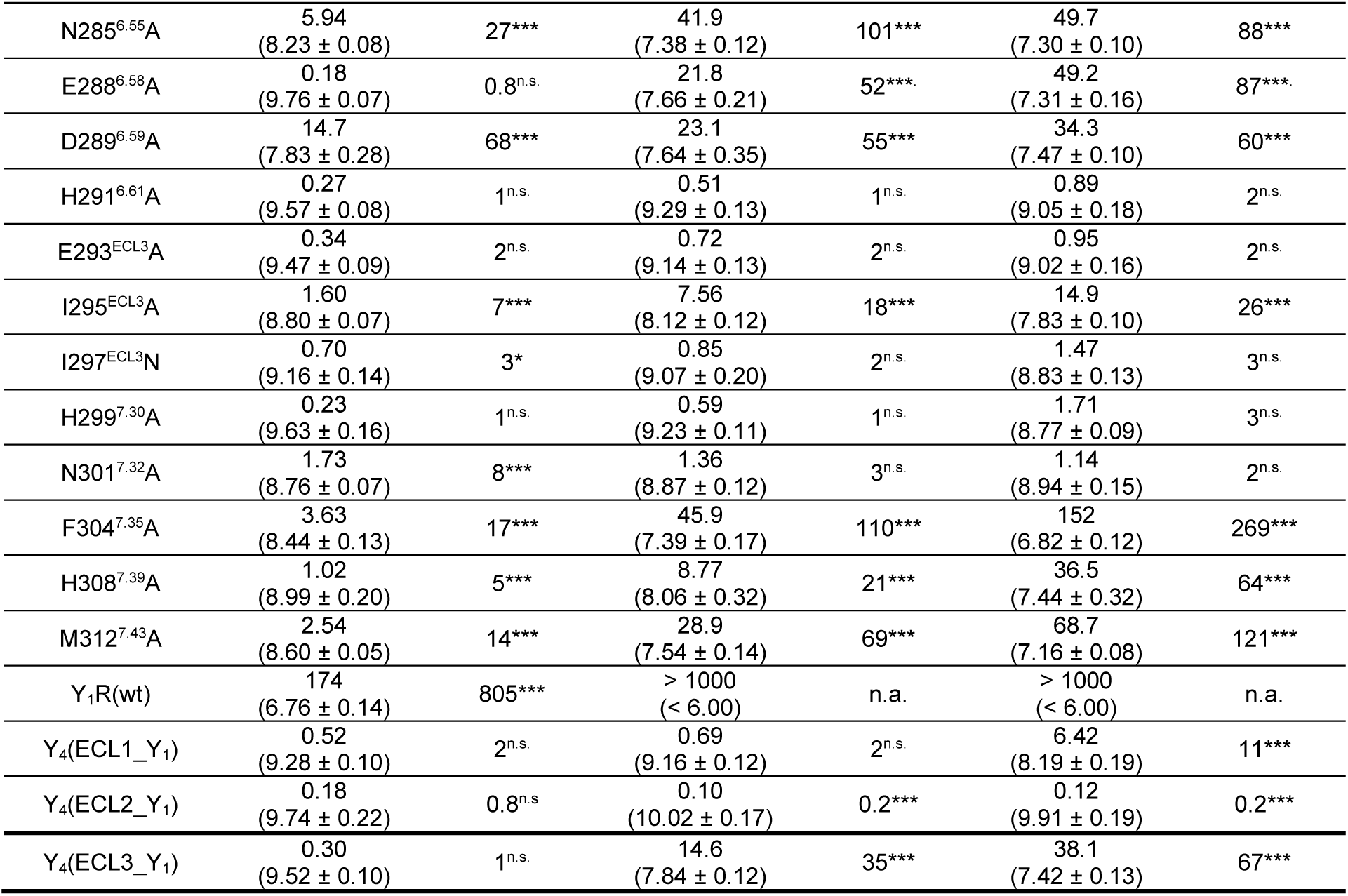
Mutagenesis studies of wild-type Y4R, Y4R single- and multiple-residue variants, wild-type Y4R, and Y4/Y1R loop chimeras with PP, UR-AK95c, and UR-AK86c. EC50 values were determined by IP-one accumulation assays in COS-7 cells transiently expressing the receptor variant-eYFP constructs and Δ6Gαqi4myr. The EC50 shift represents the ratio of the EC50 value of the peptide at the receptor variant to the EC50 value of the same peptide at wild-type Y4R. Each Y4R variant was tested in at least three independent experiments. Statistical analysis was conducted using one-way ANOVA followed by Dunnett’s post-test. Residues are named according to the Ballesteros-Weinstein nomenclature (*Ballesteros and Weinstein, 1995*). n.a., not applicable, n.s., not significant, *, P < 0.033, **, P < 0.002, ***, P < 0.001. 4xA_ECL2, Y_4_(F184A, H188A, L192A, F194A).

**Figure 3 -figure supplement 1:**
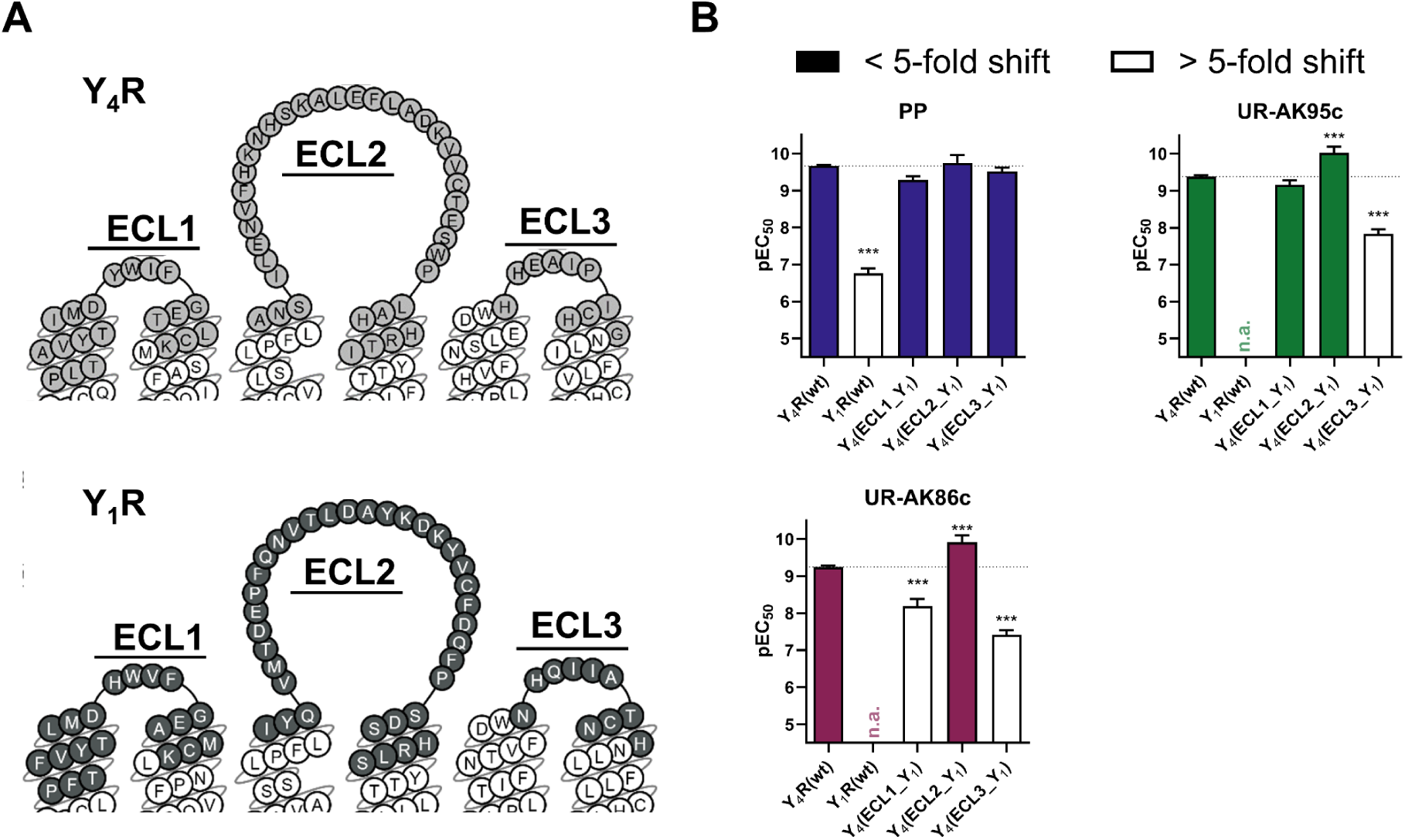
Functional studies of PP, UR-AK95c, and UR-AK86c at Y4/Y1R loop chimeras. A) Snake plots of Y4R and Y1R. Interchanged segments are colored in light grey (Y4R) and dark grey (Y1R). Snake plot was adapted from GPCRdb.org (***Pándy-Szekeres et al., 2018***). B) pEC_50_ values of PP, UR-AK95c, and UR-AK86c determined at wild-type Y4R, wild-type Y1R, and Y4/Y1R loop chimeras. Functional data were obtained through IP-one accumulation assay conducted in COS-7 cells transiently transfected with C-terminally eYFP-tagged receptor variant and Δ6Gαqi4myr. Significance was determined by one-way ANOVA with Dunnett’s post-test. *, P < 0.033, **, P < 0.002, ***, P < 0.001. All data represent the mean ± SEM from at least three independent experiments, each performed in triplicates.

**Figure 3 -figure supplement 2:**
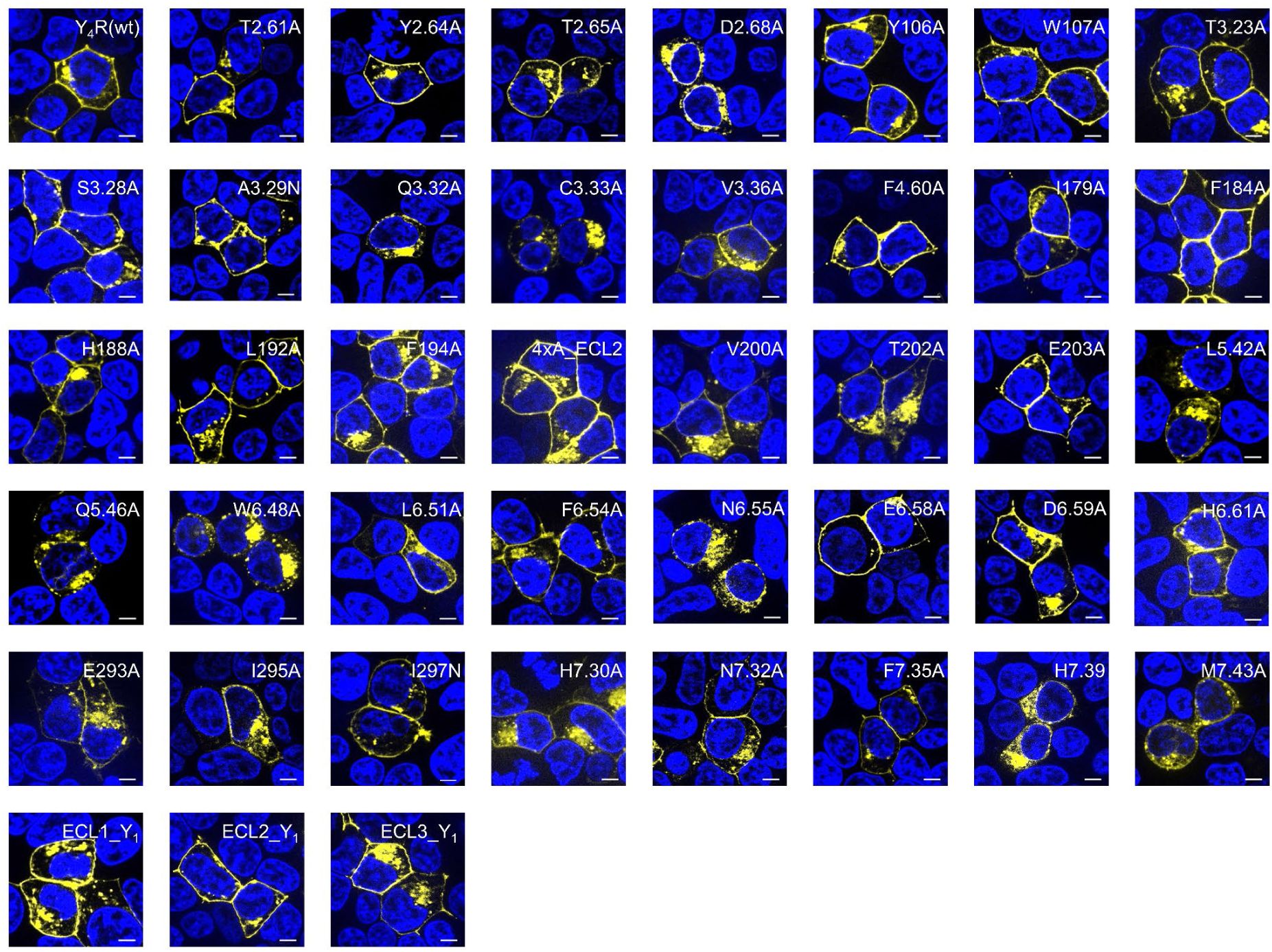
Membrane expression of Y4R residue variants and Y4/Y1R loop chimeras. The membrane insertion was studied using fluorescence microscopy in HEK293 cells, transiently expressing Y4R-eYFP fusion protein. The receptor-eYFP constructs are represented in yellow, while the nuclei stained with Hoechst33342 are shown in blue. Scale bar is 5 μm. Pictures are representatives of at least two independent experiments. 4xA_ECL2, Y_4_(F184A, H188A, L192A, F194A).

**Figure 3- figure supplement 3:**
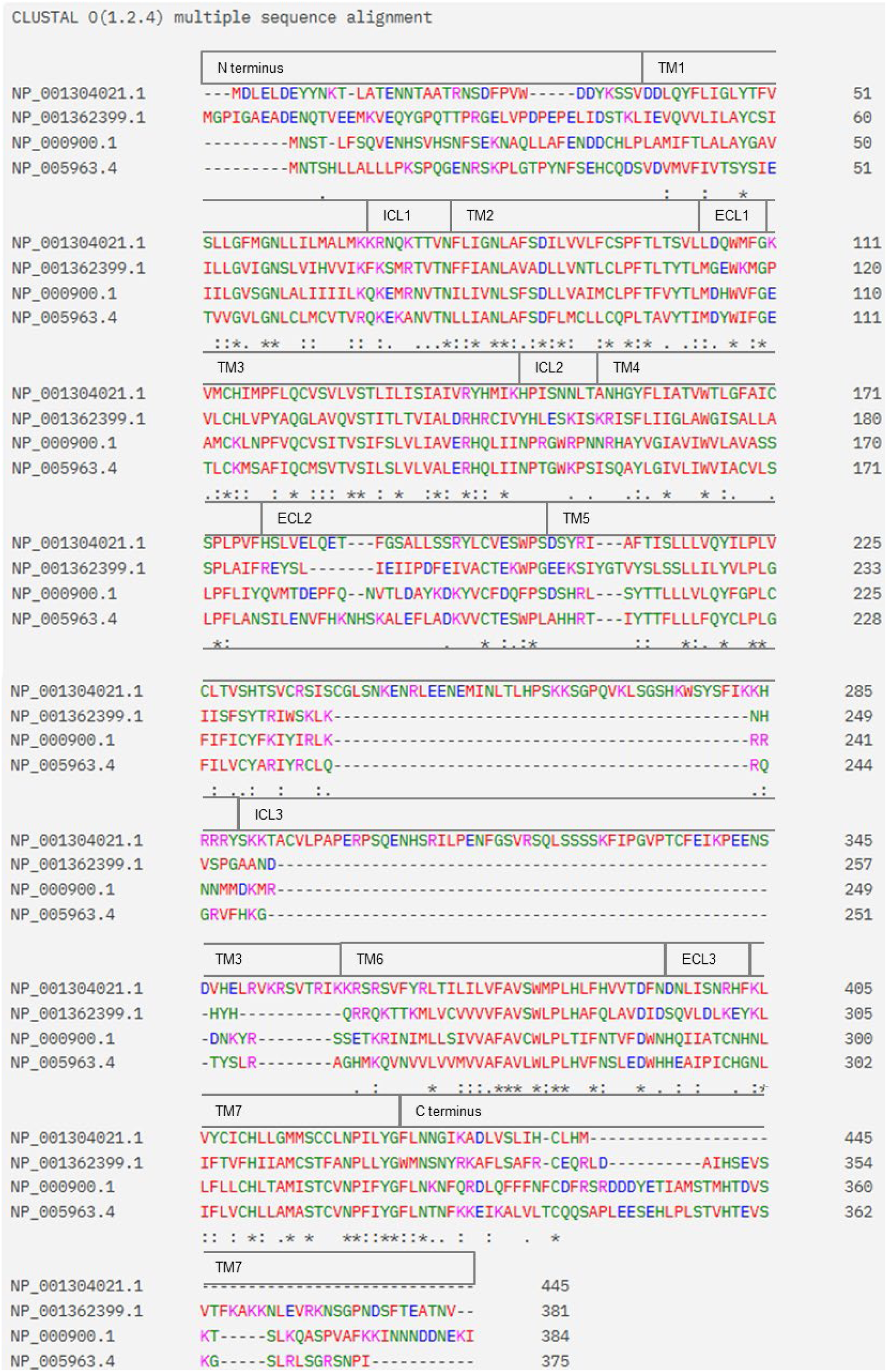
Multiple sequence alignment of human neuropeptide Y receptor subtypes. Sequences were obtained from the National Center for Biotechnology Information (NCBI) database (NIH) and represent the amino acid sequences of Y1R (NP_000900.1), Y2R (NP_001362399.1), Y4R (NP_005963.4), and Y5R (NP_001304021.1). Alignment was performed using CLUSTAL O(1.2.4) multiple sequence alignment tool (***Chenna et al., 2003***). Receptor segments are annotated based on the location of the residues in the available PP-Y4R cryo-EM structure. Residue coloring indicates side chain properties: negatively charged (blue), positively charged (pink), polar (green), and hydrophobic (red).

**Figure 5 -table supplement 1:**
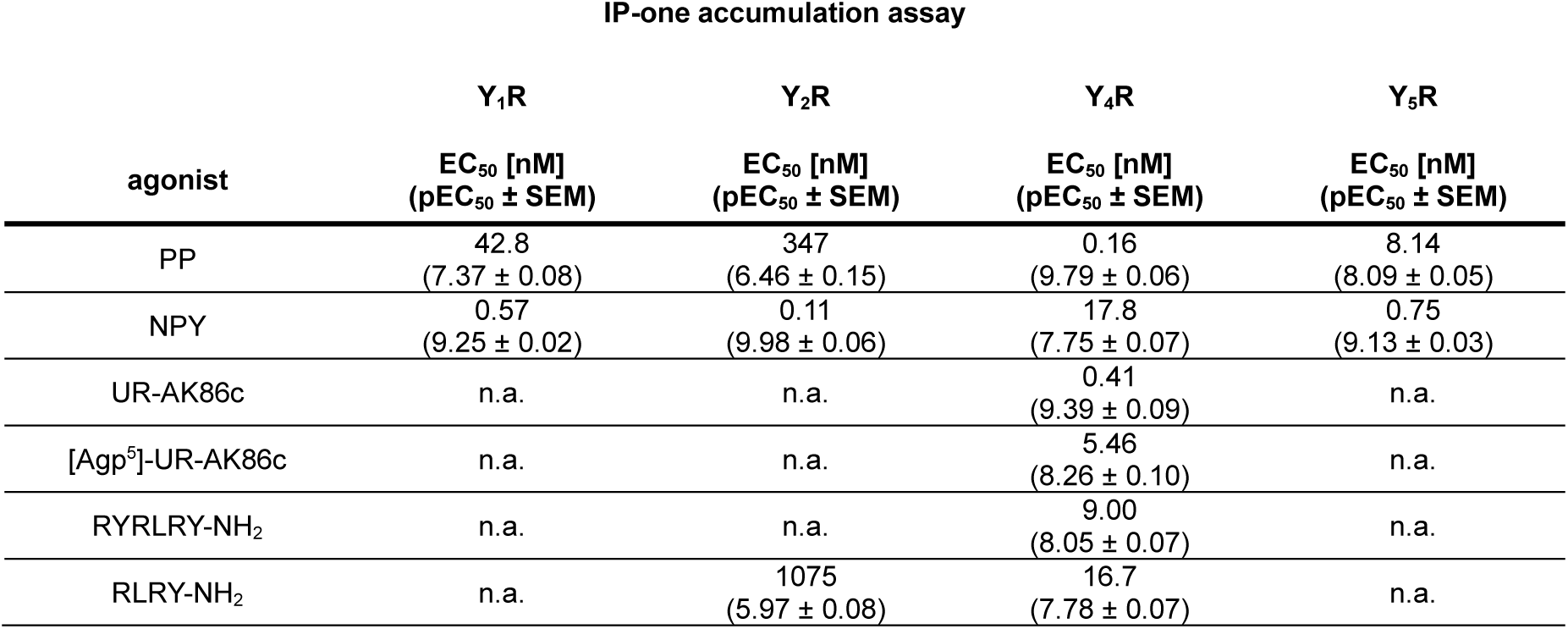
Functional data of tested peptides obtained by IP-one accumulation assays. IP-one accumulation assay was performed in COS-7 cells stably expressing the Y1/2/4/5R-eYFP construct and the chimeric G protein Δ6Gαqi4myr.. EC50 and pEC50 values were calculated from a minimum of three independent experiments. n.a., not applicable.

**Figure 5 – figure supplement 1:**
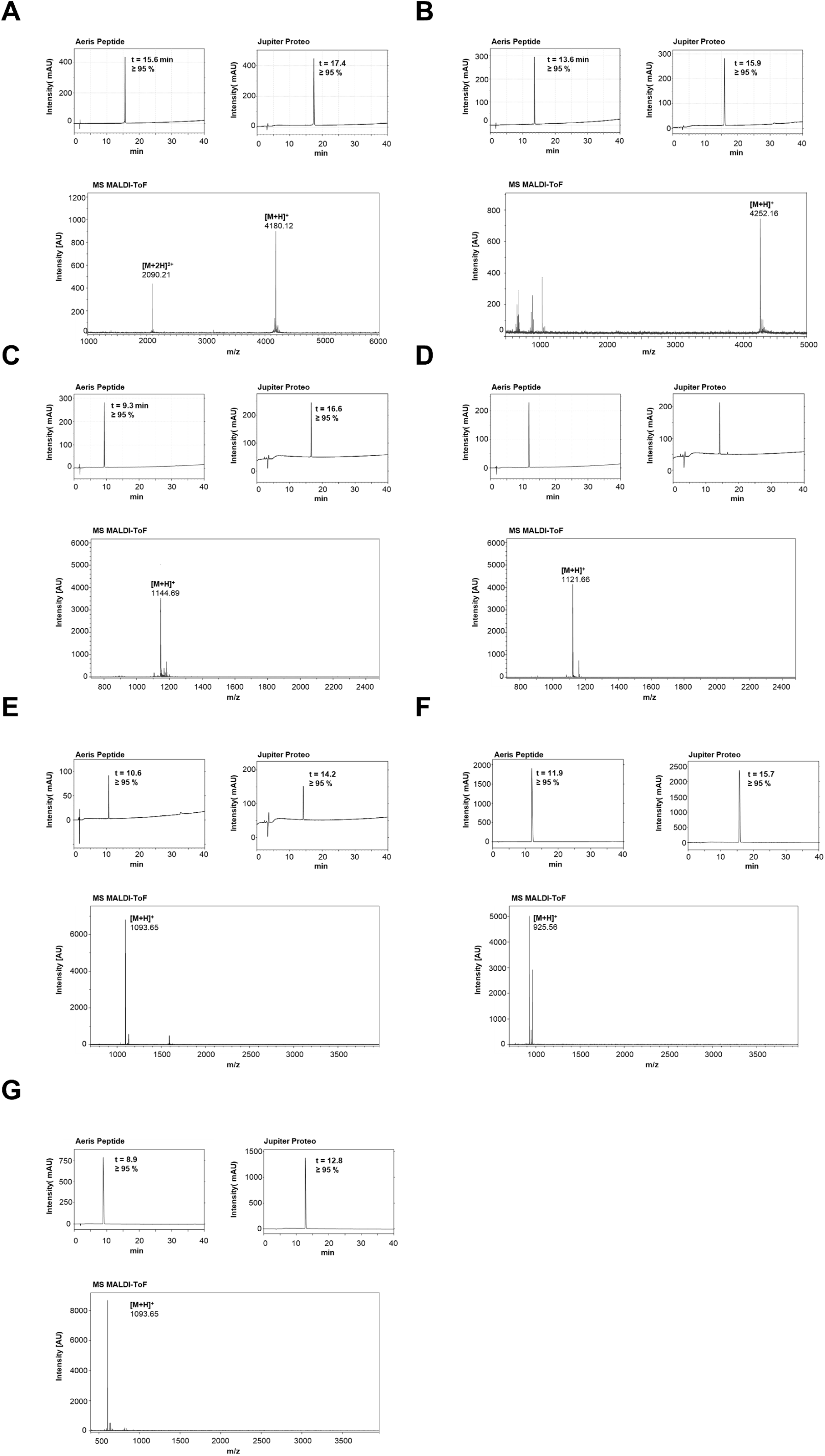
Analysis of hPP (A), pNPY (B), UR-AK95c (C), UR-AK86c (D), [Agp^5^]-UR-AK86c (E), RYRLRY-NH2 (F), RLRY-NH2 (G). The upper panels show RP-HPLC chromatograms recorded at 220 nm using gradients of 20–70% (A, B), 10–60% (C–E), or 5–55% (F, G) eluent B (0.08% TFA in ACN) in eluent A (0.1% TFA in H₂O) over 40 min at 40 °C. Analyses were performed on either an Aeris Peptide 100 Å XB-C18 column (flow rate: 1.55 mL/min) or a Jupiter Proteo 90 Å C12 column (flow rate: 1.0 mL/min). The lower panels show the corresponding MALDI-ToF mass spectra.

**Figure 6 -table supplement 1:**
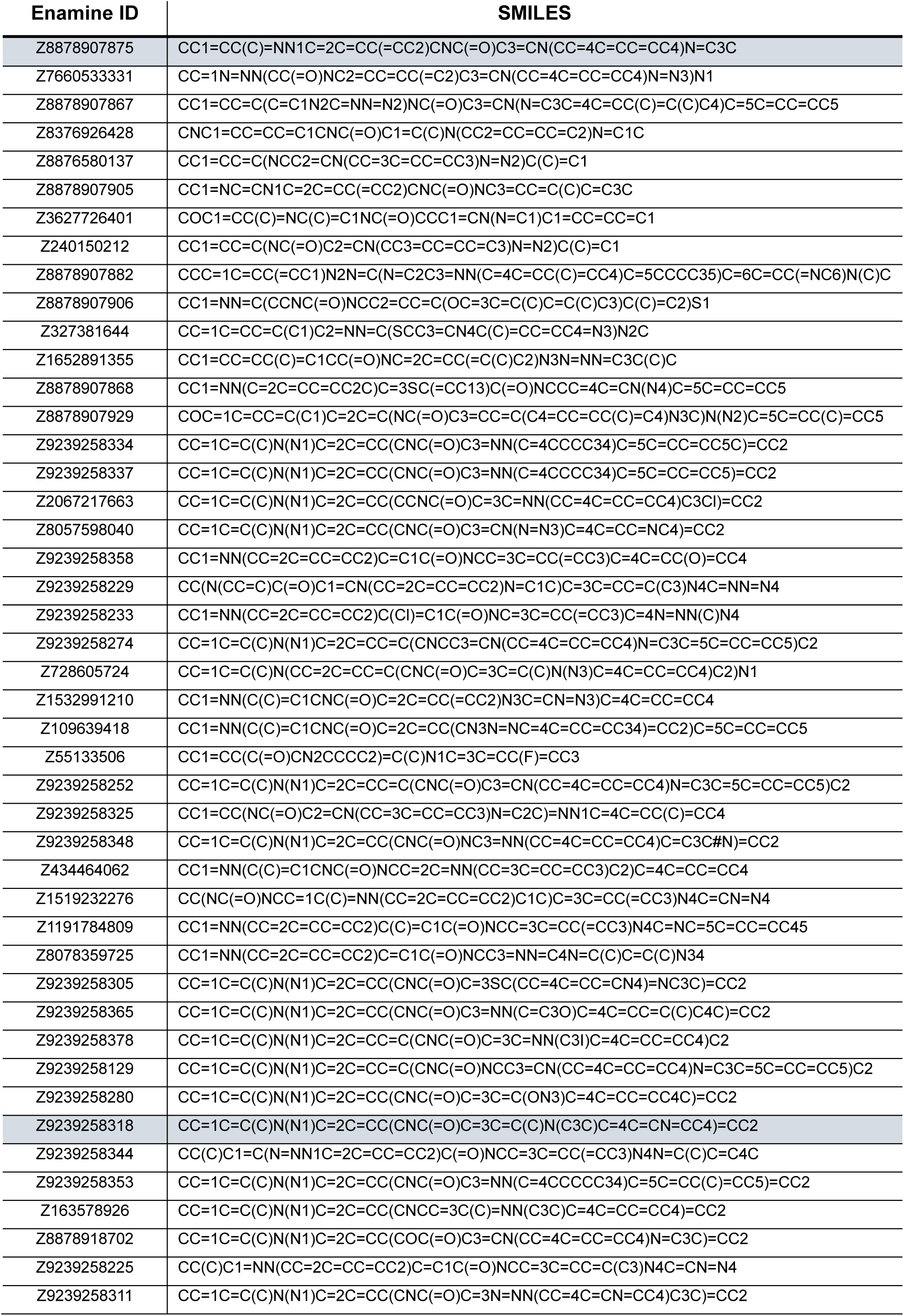
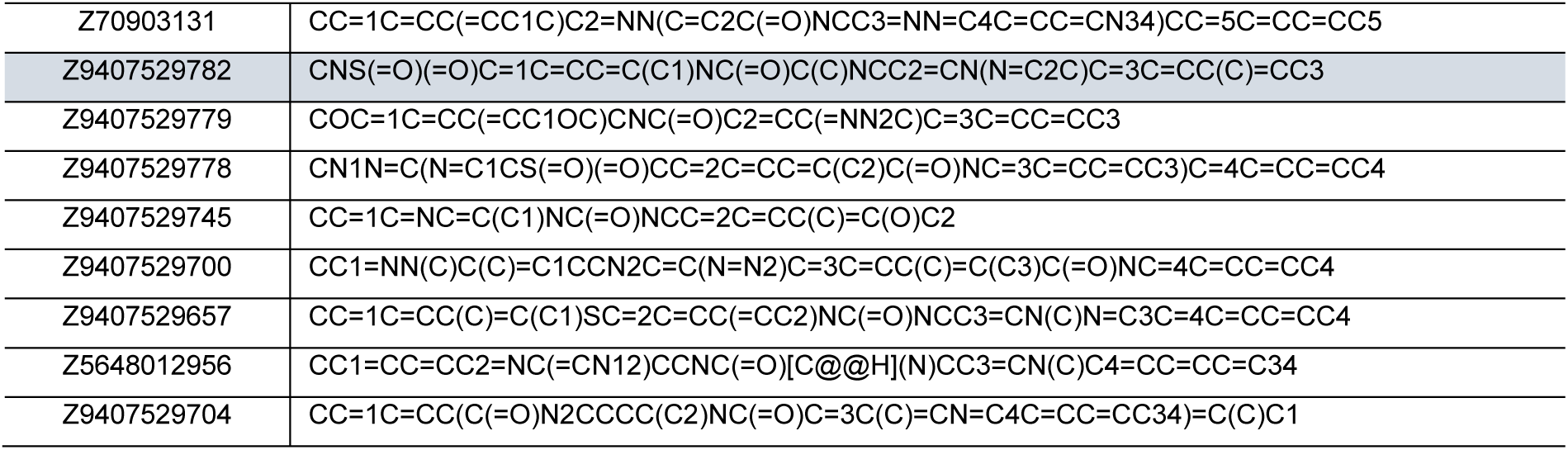
List of compounds ordered from Enamine LTD including their Enamine ID and structure as SMILES (*Grygorenko et al., 2020*). The compounds, which showed significant Y4R activity, are highlighted in grey.

## Acknowledgements

The authors gratefully thank K. Löbner, J. Schwesinger, R. Müller and C. Dammann for technical lab support. The authors appreciate the Scientific Computing Cluster Galaxy of Leipzig University for providing the computational resources. The authors would like to credit Iaroslava Kos and Enamine LTD for their support and access to their dataset, and Lukas von Bredow as a medicinal chemist for the fruitful discussions and insights into hit picking.

## Data availability

All data from this study are provided as source data files associated with the corresponding figures and figure supplements. These files are available with the article.

## Funding

**German Research Foundation (DFG, project number 421152132, SFB1423, subprojects B01, A07, Z03, Z04)**

T.P, C.S., M.S., F.L., D.J., J.S., J.M., A.G.B.-S.

**German Research Foundation (DFG, project number 460865652, SPP2363)**

F.L., V.E., P.E., J.M.

**Federal Ministry of Education and Research of Germany (BMBF, Center of Excellence for AI-research Center for Scalable Data Analytics and Artificial Intelligence (ScaDS. AI) Dresden/Leipzig, project number SCADS24B)**

F.L, P.E., J.M.

**German Research Foundation (DFG, research grant KE 1857/1-3)**

A.O.G., M.K.

**Federal Ministry of Education and Research (BMBF, German Network for Bioinformatics Infrastructure (de.NBI))**

J.M.

**German Academic Exchange Service (DAAD, School of Embedded Composite AI (SECAI 15766814))**

J.M.

**Humboldt Professorship of the Alexander Humboldt Foundation**

J.M.

**National Institute of Health (NIH, S10 OD016216, S10 OD016216, S10 OD020154)**

J.M.

### Author information

**T.P.**

- Institute of Biochemistry, Leipzig University, 04103 Leipzig, Germany
- Contribution: Conceptualization, Methodology, Investigation, Formal analysis, Data curation, Visualization, Writing—original draft
- ORCID: 0009-0004-9374-8428

#### C.S

- Institute of Biochemistry, Leipzig University, 04103 Leipzig, Germany
- Contribution: Conceptualization, Methodology, Investigation, Formal analysis, Data curation, Writing—review & editing
- ORCID: 0000-0002-2689-0609

**M.S.**

- Institute for Drug Discovery, Leipzig University, 04103 Leipzig, Germany
- Contribution: Methodology, Investigation, Software, Writing—review & editing
- ORCID: 0009-0001-3939-8138

**F.L.**

- Institute for Drug Discovery, Leipzig University, 04103 Leipzig, Germany, AI-Driven Therapeutics GmbH, Humboldtstr. 25, 04105 Leipzig, Germany
- Contribution: Methodology, Investigation, Software, Writing—review & editing
- ORCID: 0009-0007-9199-1629
- Competing interest: The author declares that he has no known competing financial interests or personal relationships that could have appeared to influence the work reported in this paper. AI-Driven Therapeutics was not financially involved in the study.

**D.J.**

- Institute for Drug Discovery, Leipzig University, 04103 Leipzig, Germany
- Contribution: Methodology, Investigation, Software, Writing—review & editing
- ORCID: 0009-0004-7535-9314

**V.E.**

- Institute for Drug Discovery, Leipzig University, 04103 Leipzig, Germany
- Contribution: Methodology, Investigation, Software
- ORCID: 0009-0000-7382-0147

**P.E.**

- Institute for Drug Discovery, Leipzig University, 04103 Leipzig, Germany, AI-Driven Therapeutics GmbH, Humboldtstr. 25, 04105 Leipzig, Germany
- Contribution: Methodology, Investigation, Software, Writing—review & editing
- ORCID: 0009-0006-7379-5096
- Competing interest: The author declares that he has no known competing financial interests or personal relationships that could have appeared to influence the work reported in this paper. AI-Driven Therapeutics was not financially involved in the study.

**J.S.**

- Institute of Biochemistry, Leipzig University, 04103 Leipzig, Germany
- Contribution: Methodology, Investigation
- ORCID: 0009-0002-4633-7656

**A.O.G.**

- Institute of Pharmacy, University of Regensburg, 93040 Regensburg, Germany
- Contribution: Investigation, Resources, Writing—review & editing
- ORCID: 0000-0001-5166-2168

**M.K.**

- Institute of Pharmacy, University of Regensburg, 93040 Regensburg, Germany
- Contribution: Resources, Writing—review & editing
- ORCID: 0000-0002-8095-8627

**J.M.**

- Institute for Drug Discovery, Leipzig University, 04103 Leipzig, Germany; Center for Structural Biology, Vanderbilt University, Nashville, TN 37240, USA; Department of Chemistry, Department of Pharmacology and Institute of Chemical Biology, Vanderbilt University, Nashville, TN 37235, USA; School of Embedded Composite Artificial Intelligence SECAI, 04105 Leipzig, Germany; Faculty of Mathematics and Informatics, Faculty of Chemistry, Leipzig University, 04103 Leipzig, Germany
- Contribution: Conceptualization, Supervision, Project administration, Funding acquisition, Resources, Writing—review & editing
- ORCID: 0000-0001-8945-193X

**A.G.B.-S.**

- Institute of Biochemistry, Leipzig University, 04103 Leipzig, Germany
- Contribution: Conceptualization, Supervision, Project administration, Funding acquisition, Resources, Writing—review & editing
- ORCID: 0000-0003-4560-8020

## Notes

### Competing Interest Statement

The authors have declared no competing interest.

